# Three-dimensional genomic mapping of human pancreatic tissue reveals striking multifocality and genetic heterogeneity in precancerous lesions

**DOI:** 10.1101/2023.01.27.525553

**Authors:** Alicia M Braxton, Ashley L Kiemen, Mia P Grahn, André Forjaz, Jaanvi Mahesh Babu, Lily Zheng, Liping Jiang, Haixia Cheng, Qianqian Song, Rebecca Reichel, Sarah Graham, Alexander I Damanakis, Catherine G Fischer, Stephanie Mou, Cameron Metz, Julie Granger, Xiao-Ding Liu, Niklas Bachmann, Cristina Almagro-Pérez, Ann Chenyu Jiang, Jeonghyun Yoo, Bridgette Kim, Scott Du, Eli Foster, Jocelyn Y Hsu, Paula Andreu Rivera, Linda C Chu, Fengze Liu, Noushin Niknafs, Elliot K Fishman, Alan Yuille, Nicholas J Roberts, Elizabeth D Thompson, Robert B Scharpf, Toby C Cornish, Yuchen Jiao, Rachel Karchin, Ralph H Hruban, Pei-Hsun Wu, Denis Wirtz, Laura D Wood

**Author notes:** Authors contributed equally (Alicia M Braxton, Ashley L Kiemen). To whom correspondence should be addressed: Laura D. Wood, MD, PhD, CRB2 Room 345, 1550 Orleans Street, Baltimore, MD 21231, Denis Wirtz, PhD, 265 Garland Hall, 3400 N. Charles Street, Baltimore, MD 21218, Yuchen Jiao, PhD, 4127 Laobingfang Building, 17 Panjiayuannali, Cancer Hospital, Chinese Academy of Medical Sciences, Beijing, China, 86-10-87787615.

## Abstract

Pancreatic intraepithelial neoplasia (PanIN) is a precursor to pancreatic cancer and represents a critical opportunity for cancer interception. However, the number, size, shape, and connectivity of PanINs in human pancreatic tissue samples are largely unknown. In this study, we quantitatively assessed human PanINs using CODA, a novel machine-learning pipeline for 3D image analysis that generates quantifiable models of large pieces of human pancreas with single-cell resolution. Using a cohort of 38 large slabs of grossly normal human pancreas from surgical resection specimens, we identified striking multifocality of PanINs, with a mean burden of 13 spatially separate PanINs per cm^3^ of sampled tissue. Extrapolating this burden to the entire pancreas suggested a median of approximately 1000 PanINs in an entire pancreas. In order to better understand the clonal relationships within and between PanINs, we developed a pipeline for CODA-guided multi-region genomic analysis of PanINs, including targeted and whole exome sequencing. Multi-region assessment of 37 PanINs from eight additional human pancreatic tissue slabs revealed that almost all PanINs contained hotspot mutations in the oncogene *KRAS*, but no gene other than *KRAS* was altered in more than 20% of the analyzed PanINs. PanINs contained a mean of 13 somatic mutations per region when analyzed by whole exome sequencing. The majority of analyzed PanINs originated from independent clonal events, with distinct somatic mutation profiles between PanINs in the same tissue slab. A subset of the analyzed PanINs contained multiple *KRAS* mutations, suggesting a polyclonal origin even in PanINs that are contiguous by rigorous 3D assessment. This study leverages a novel 3D genomic mapping approach to describe, for the first time, the spatial and genetic multifocality of human PanINs, providing important insights into the initiation and progression of pancreatic neoplasia.

## INTRODUCTION

Pancreatic ductal adenocarcinoma (PDAC) is an aggressive malignancy that is frequently diagnosed at an advanced stage, leading to a five-year survival rate of just 11%.^1,2^ However, PDAC arises from noninvasive precursor lesions, the most common of which is pancreatic intraepithelial neoplasia (PanIN), which are curable if detected and treated early.^3^ PanIN lesions are challenging to study; due to their small size (by definition <0.5cm in a standard histological section), PanINs cannot be identified from gross examination of pancreatic tissue and are only visible via microscopic examination of tissue sections.^4,5^ To date, studies of PanINs in human pancreatic tissue samples have evaluated discrete histological sections.^6–9^ While this approach may identify portions of PanIN lesions, such two-dimensional (2D) analyses cannot accurately assess the number, size, shape, or connectivity of PanINs in a tissue sample, key features to understand the earliest steps of pancreatic tumorigenesis. A three-dimensional (3D) analysis of larger slabs of tissue is needed to assess these features more completely. We recently reported CODA, a novel machine-learning pipeline for 3D image analysis that can be used to generate quantifiable models of large pieces of human pancreas with single-cell resolution.^10^ Using CODA, we can now quantify the number and connectivity of PanINs through systematic evaluation of large pieces of human pancreatic tissue.

The accumulation of somatic mutations in oncogenes and tumor suppressor genes drives the initiation and progression of PanINs.^3^ These include hotspot mutations in the oncogene *KRAS*, which are thought to initiate pancreatic ductal neoplasia and occur in more than 90% of invasive pancreatic cancers, as well as inactivating mutations in tumor suppressor genes, including *CDKN2A* and *TP53*, which occur at lower prevalence.^3,6,11–14^ Previous studies have reported heterogeneity in driver gene mutations in intraductal papillary mucinous neoplasms, large non-invasive cystic precursors to PDAC, using multi-region next generation sequencing, demonstrating the complex clonal evolution in these lesions.^15–17^ When examined, such heterogeneity in driver gene mutations has not been identified in primary PDACs or metastases, but the presence of driver gene heterogeneity in PanINs has not been assessed.^18,19^ Genetic analyses guided by CODA models allows exhaustive assessments of the genetic heterogeneity in PanINs, as spatially separate PanIN lesions can be identified in 3D and then separately sequenced.

This study comprehensively describes the 3D microanatomy, cellular features, and somatic genetic alterations of human PanINs. We generated 3D models from a large cohort of grossly normal surgically resected human pancreatic tissue slabs using CODA. In a subset of slabs, we created a novel workflow for mapping genetic variation across 3D structures through integration of CODA-generated 3D microanatomy with multi-region targeted and whole exome sequencing. With this workflow, we assessed both intra-PanIN and inter-PanIN genetic heterogeneity. Together, these data reveal striking multifocality of PanINs and elucidate their complex molecular origins.

## METHODS

### Specimen Acquisition

This study was approved by the Institutional Review Board at The Johns Hopkins Hospital. Thick slabs of grossly normal pancreatic tissue were harvested from surgical pancreatectomy specimens and consecutively assigned a slab number. For 3D modeling, a cohort was selected to investigate PanINs arising in otherwise histologically normal blocks of pancreatic parenchyma. Following histologic review by an expert pancreatic pathologist, slabs containing significant fibrosis, atrophy, or PDAC in the analyzed tissue slab were excluded. In addition, slabs with a diagnosis of intraductal papillary mucinous neoplasm (IPMN) were excluded due to the histological similarities between PanIN and IPMN and the propensity of IPMN to involve smaller ducts, complicating the reliable distinction of the two precursor lesions.^5^ With these criteria, we selected 38 slabs of pancreas tissue from 38 patients for 3D anatomical modeling. Twelve of these patients had PDAC elsewhere in their pancreas. The remaining 26 patients had other pancreatic neoplasms elsewhere in their resection specimens, including pancreatic neuroendocrine tumors, serous cystadenomas, distal common bile duct adenocarcinomas, metastatic carcinomas from other organ sites, mucinous cystic neoplasms, tubulovillous adenomas of the duodenum, ampullary tumors, and lymphoepithelial cysts (Supplementary Table 1).

A separate cohort of grossly normal pancreatic tissue slabs underwent combined anatomic and genomic analyses. Because of the unique sectioning scheme (details below) used for 3D modeling and next generation sequencing (NGS), which included a large number of specialized membrane slides for microdissection, criteria for inclusion were based on the first histologic slide cut from the surface of each slab for this cohort, which was assessed prior to full sectioning. The criteria were as follows: PanIN present, PDAC and significant fibrosis absent on the first slide sectioned. Any patients with a clinical diagnosis of IPMN were excluded. Using these criteria, eight additional slabs of tissue were selected from eight patients for 3D modeling and NGS – these patients did not overlap with the 38 patients analyzed by anatomic modeling only. Three of these eight patients had a pathological diagnosis of PDAC elsewhere in their pancreas. The remaining five of the eight patients had neoplasms not affecting the pancreatic ductal system, including well-differentiated pancreatic neuroendocrine tumors (2), serous cystadenoma (1), distal common bile duct adenocarcinoma (1), and colon cancer metastatic to the pancreas (1).

### Sample processing

Each of the 46 slabs of harvested tissue were formalin-fixed and paraffin-embedded, followed by complete serial sectioning at 5μm. Every third slide was stained with H&E and digitized at 20x magnification for 3D modeling. As previously described, skipping staining on two out of every three slides does not lead to any significant loss in micro-anatomical information. ^10^ Sample processing yielded a mean of 1,288 slides per block (range 679-1,703) and mean tissue volume of 2.03 cm^3^ (range 0.94-3.62cm^3^). All PanIN lesions present on every 50^th^ slide were manually annotated using Aperio ImageScope to verify accuracy of the generated 3D models.^10^ For the eight slabs undergoing NGS, every third slide was cut onto membrane slides (Zeiss Membrane Slide 1.0 PEN; Carl Zeiss, Oberkochen, Germany) for laser capture microdissection (LCM).

### 3D reconstruction of serially sectioned H&E sections of human pancreas

Using the previously validated method CODA, we converted the serially sectioned histological slides of human pancreas tissue into digital 3D maps of pancreas microanatomy for all 46 slabs.^10^ Briefly, the CODA workflow can be split into four steps: image registration, single-cell detection, tissue segmentation, and 3D visualization. For a pair of images, the registration maximizes the 2D cross correlation of pixel intensity to align all images and correct for tissue rotation, translation, folding, splitting, and stretching. A cell detection algorithm is used to quantify the cellularity of components via detection of 2D intensity peaks in the hematoxylin channel of the H&E images. Deep learning is next used to create microanatomical labels from the histological images. The trained algorithm was used to label, to a resolution of 2 μm, eight microanatomical structures in histological images of the pancreas: islets of Langerhans, normal ductal epithelium, vasculature, fat, acinar tissue, collagen, PanIN, and PDAC. Samples containing regions of cancer or lymph nodes were labelled using a separate model trained to additionally recognize these components. The image registration, cell detection, and tissue segmentation are integrated to create 3D reconstructions of pancreas microanatomy at large scale (up to multi-cm^3^), while maintaining single-cell resolution.

All 46 thick slabs of pancreas tissue were reconstructed using CODA. The 3D datasets used for quantification and visualizations were subsampled from the classified resolution of 2 micron per pixel per image to an isometric resolution of 12 x 12 x 12 μm. The cell detection algorithm was manually validated on four regions of randomly selected images and cell detection parameters were adjusted until a minimum accuracy of 90% was obtained for each tissue component. For the deep learning, all available annotated images were collected: 90% of images were used for training and 10% of images were used as an independent testing set of model accuracy across unseen images in the analyzed cohort. Models were deemed acceptable when they achieved >90% per class precision and recall on the independent testing images.

### Validation of PanIN detection in samples

All objects labelled as PanIN by the segmentation algorithm were post-processed to ensure that no non-neoplastic regions were counted as PanIN. In the 3D volume matrix saved at a resolution of 12 x 12 x 12 μm^3^, objects of fewer than 20 voxels and objects present on fewer than three histological sections were eliminated. The histology of all remaining objects was manually assessed to determine whether they were PanIN. The 3D bounding box of each object was determined. The regions of the serial H&E images contained in this bounding box were extracted from the registered 5x magnification image stack and were saved as a separate image stack (Supplementary Video 1). These stacks were manually viewed using FIJI ImageJ.^20^ A matrix was created in Microsoft excel containing the PanIN identifier (labeled from A to Z based on PanIN), and the true determined label of the region. All regions determined to be false positive PanIN labeling during manual validation were corrected in the 3D matrices. False negatives were assessed through comparison of the deep learning labelled structures to manual pathologist-guided annotations of neoplastic tissue on one in every fifty sections throughout the samples.

### Color-coded labelling of connected regions of PanIN in 2D H&E images

For the eight slabs that were assessed by the combined 3D anatomic and genomic analyses, videos with exhaustive, color-coded annotations of PanIN lesions were created using the 3D tissue models so that connected regions of each PanIN could be efficiently identified, microdissected, and collected in separate vials for genomic analysis. For this purpose, PanIN labels were collected from the digital 3D tissue matrix. Independent (i.e. discontinuous in 3D) PanINs were sorted from largest to smallest, and the ten largest PanIN lesions were assigned distinct colors (largest PanIN assigned pink, second largest PanIN assigned light blue, third largest PanIN assigned yellow, etc.) for easy visualization. For samples with more than ten identified PanIN lesions, the eleventh through the smallest were assigned the same color (olive green), as these PanIN were too small to be microdissected and sequenced. Next, we determined which PanIN lesions were present in each z-plane of the 3D structure. For each plane, the 5x registered H&E image was loaded into MATLAB 2021b. An outline of the regions of each PanIN present in that plane was digitally thickened and was overlayed on the H&E image in the correct location and in the color assigned to that PanIN. This process created an H&E image with color outlines highlighting all PanIN lesions in the section, and further highlighting any connectivity, and size order (via color). These 5x images were saved separately as jpg files, and together as a lower-resolution stacked video (Supplementary Video S2).

### Calculation of 3D structural features of PanIN lesions

The 3D digital tissue matrices that defined tissue types and cell coordinates were used to calculate a range of tissue properties. Each PanIN lesion was separately identified using MATLAB 2021b. By summing the number of voxels in each separate PanIN lesion (and converting from voxel to mm3), the volume of each PanIN lesion was obtained. The volume of a PanIN was defined as the volume of the neoplastic cells, and excluding the lumina within the glands. The Regionprops3 command in MATLAB 2021b was used to determine the length of the primary axes containing each PanIN. Through dot multiplication of the PanIN ID matrix with the matrix containing the cell coordinates, the number of cells in each PanIN could be calculated. Cell counts generated from the 2D histological sections were extrapolated to 3D space using a previously developed technique based on cell-type-dependent nuclear measurements, as well as the thickness and spacing of the histological slides.^10^

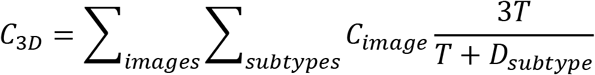

Finally, the cell density of each PanIN lesion was determined by dividing the number of cells of each lesion by its volume (excluding the lumen). The total volume, cellularity, and cell density of all other tissue components were similarly calculated using the 3D tissue and cell matrices.

### Laser capture microdissection

For NGS, membrane slides from 8 of the 46 tissue slabs were deparaffinized in xylenes for five minutes, and dehydrated with 100%, 95%, and 70% ethanol for one minute each. Slides were stained with crystal-violet (Sigma Aldrich; diluted 1:4 in 70% ethanol) for 30 seconds and dehydrated by ascending ethanol solutions. The stained slides were microdissected the same day. For genetic analyses assessing intra- and inter-PanIN heterogeneity, each slab was divided into five regions of equal size along the z-axis. On average, tissue slabs spanned 778 slides (range 500-1,000) and each region covered 146 slides (range 100-200) (Supplementary Table 3). Each spatially unconnected PanIN lesion identified in the corresponding 3D models was microdissected (Leica LMD7000 instrument; Wetzlar, Germany) into a separate collection tube (autoclaved 0.5mL Qubit Assay Tubes, Invitrogen; Waltham, MA) for each region. Spatially distinct PanIN lesions and PDAC on H&E images were annotated by CODA utilizing colors corresponding to 3D models; these annotated images were used to guide microdissection of distinct lesions into separate collection tubes. LG and HG components of PanIN lesions were isolated separately by microdissection, as was any PDAC identified in the tissue. Matched control samples from each patient were obtained from clinical archives for sequencing, including matched normal tissue (duodenum or spleen) and matched PDAC. For matched control samples, one 5μm section was cut onto a regular slide, deparaffinized as described above, and scraped into a collection tube utilizing a sterile razor blade.

### DNA Extraction and Quantification

DNA was extracted using the QIAamp DNA FFPE Tissue Kit (Qiagen; Hilden, Germany) following manufacturer’s recommendations with the following modifications: after overnight digestion, samples were sheared to 200-350 base pair fragments using the Covaris S220 Sonicator (Covaris; Woburn, MA). Subsequently, 10uL of proteinase K was added and samples were digested for one hour at 56°C before resuming manufacturer provided protocol. DNA quantity and fragment size were quantified using an Agilent Bioanalyzer according to manufacturer’s recommendations. Sample processing was the same for matched normal tissue and matched PDAC tissue. Samples were stored at −20°C until library preparation.

### Library Preparation and Targeted Sequencing

All PanIN regions that yielded at least 10ng of DNA in the eight slabs used for combined 3D anatomic and genomic analysis were assessed by targeted next generation sequencing using a custom panel. The commercially available ClearSeq Comprehensive Cancer (Agilent; Santa Clara, CA) targeted panel was selected. Using the Agilent SureDesign software, we customized the panel to induce baits for the coding regions of additional genes important in pancreatic cancer development. In total, the panel covered 154 well-characterized cancer driver genes, including all major drivers of pancreatic ductal neoplasia (Supplementary Table 3). 10-200ng of DNA was utilized per sample for library preparation, following the manufacturer’s protocol (SureSelect XT HS2 DNA kit; Agilent). Barcoded individual samples were pooled following the manufacturer’s recommendations and sequenced on a MiSeq (Illumina, San Diego, CA) generating 2 x 150 base-paired reads.

### Targeted Sequencing Analysis

Sequencing data were processed by following “The Genome Analysis Toolkit (GATK) Best Practices Pipeline” ^21^. FASTQ reads were converted to an unmapped BAM file and adapter sequences were marked. FASTQ reads were aligned to the human reference genome hg38 using Burrows-Wheeler aligner (BWA) MEM software and merged with the unmapped BAM file ^22^. The raw mapped reads were then marked for duplicate reads and underwent base quality score recalibration (GATK version v4.2.0.0). Mutect2 was used to call somatic single nucleotide variants (SNVs) and insertions/deletions (indels) ^23^. A panel of normals (PON) was created using targeted sequencing data from the eight matched normal samples. Next, Mutect2 was run for each tumor sample with its matched normal and the PON using default settings. All somatic mutations were annotated with OpenCRAVAT (version 2.2.1). Somatic mutations were subsequently filtered with the following criteria: tumor sample coverage ≥ 15X; normal sample coverage ≥ 10X; tumor frequency ≥ 0.05; ≥ 5 distinct reads supporting the mutation in tumor sample; normal frequency < 0.05. Variants were filtered to remove all noncoding variants and variants present in any normal sample, retaining coding SNVs and indels and splice site variants. Mutations meeting these criteria were used for downstream analysis. Candidate mutations were confirmed or rejected via visual inspection in Integrated Genome Viewer (IGV) ^24^. The positions of *KRAS* hotspot mutations (codons 12, 13, and 61) were visually inspected in all samples and included if VAF ≥1% and ≥ 3 distinct reads supporting the mutation.

### Whole Exome Sequencing and Analysis

PanIN regions with sufficient DNA in the eight slabs used for combined 3D anatomic and genomic analysis were also analyzed by whole exome sequencing in addition to the targeted sequencing described above. Mutation Capsule (MC) technology was applied to profile a FFPE DNA sample with both hybridization-based exome sequencing and amplification-based deep sequencing, as recently described^25^. Approximately 200ng of the previously fragmented DNA was subjected to end-repair, dA-tailing, and ligation to a customized adapter with random DNA barcodes as a unique identifier (UID) tag ^26^. The ligation product was amplified through 10 reaction cycles to generate a whole genome library (MC library) using NEBNext Ultra II DNA library Prep Kit for Illumina (New England Biolabs) and 750ng MC library was used for exome enrichment. The exome region of the whole genome libraries was enriched with the Agilent SureSelectXT Human All Exon Kit V6 (Agilent; Santa Clara, CA) and sequenced on the NovaSeq 6000 Sequencing System (Illumina) with 2 × 150-bp paired-end reads. FASTQ files were preprocessed to remove the adapter sequences ^26^. The low-quality reads were subsequently removed using Trimmomatic (v0.36). FASTQ reads were aligned to the human reference genome hg38 using Burrows-Wheeler aligner (BWA) MEM software for both tumor and matched normal samples. Mutect2 was used to call somatic variants between the tumor-normal pairs, as well as the tumor only samples against a PON, utilizing default parameters ^23^. The somatic mutations were filtered according to the following criteria: (1) the variant coverage in tumor sample was ≥ 15X, the variant allele frequency ≥ 0.10 and ≥ 7 distinct reads supporting the mutation in tumor sample; (2) the corresponding variant coverage in normal sample ≥ 10X, the variant allele frequency < 0.05. Then, all somatic mutations were annotated with vcf2maf-1.6.19. Noncoding variants and variants present in matched normal were filtered, retaining coding SNVs and indels and splice site variants. Mutations meeting these criteria were used for downstream analysis. Candidate mutations were confirmed or rejected via visual inspection in Integrated Genome Viewer (IGV) ^24^. The positions of *KRAS* hotspot mutations (codons 12, 13, and 61) were visually inspected in all samples and included if VAF ≥1% and ≥ 3 distinct reads supporting the mutation.

### Targeted sequencing of *KRAS* mutations

Ultra-deep sequencing of the *KRAS* hotspot positions was performed on the PanIN regions analyzed by whole exome sequencing. 200 ng MC library was used to profile *KRAS* mutation with deep sequencing, as previously described ^26^. Briefly, the target regions were amplified together with the DNA barcode (UID) in the adapter of the MC library for 9 cycles using a target-specific primer and a primer matching the universal sequence in the adapter. A second round of 14 cycles of PCR with one pair of nested primers matching the adapter and the target region was used to further enrich the target region and add the Illumina sequencing adapter. The amplified libraries were sequenced on the NovaSeq 6000 Sequencing System (Illumina) using 2 × 150-bp paired-end reads. The target regions were analyzed to confirm the mutation status as previously described ^26^. Briefly, the FASTQ file were preprocessed to extract UID tags ^26–28^. The residual Illumina adapters sequences and low-quality reads were subsequently removed using Trimmomatic (v0.36). The cleaned reads were mapped to the human reference genome hg19 using BWA software (BWA, v0.7.15) ^27^. BAM files were locally realigned and the base quality scores were recalibrated using Genome Analysis Toolkit (GATK, v3.1). The ‘mpileup’ command in SAMtools (v 0.1.16) was used to identify SNVs and indels ^29^. To ensure accuracy, the reads with the same UID tag were grouped into a UID family. If > 80% of reads in a UID family harbored the same variant and it contained at least two reads, the UID family was defined as an effective unique identifier family (EUID family). The prevalence of each mutation was calculated by dividing the number of mutant EUID families by the total number of the mutant and wild type EUID families. Candidate variants were annotated with the VEP (v83) and Oncotator ^30,31^. The criteria we adopted for retaining a somatic mutation was that it had an allele fraction of ≥ 1% and ≥ 7 UID. The retained mutations were verified manually using IGV ^24^.

### Pancreas Computed Tomography

An independent group of 807 individuals who were candidates for renal donation were scanned as a part of their routine care. These images were obtained. Patients were scanned on a dual-source Multidetector Computed Tomography (MDCT) scanner (Somatom Definition, Somatom Defi-nition Flash, or Somatom Force, Siemens Healthineers), or a 64-MDCT scanner (Somatom Sensation 64, Siemens Healthineers). Patients were injected with 100–120 mL iohexol (Omnipaque, GE Healthcare) at an injection rate of 4–5 mL/s. Scan protocols were customized for each patient to minimize dose and included a tube voltage of 100–120 kVp, effective tube current-exposure time product of 250–300 mAs, and pitch of 0.6–0.8. The collimation was 128 × 0.6 mm or 192 × 0.6 mm for the dual-source scanner and 64 × 0.6 mm for the 64-MDCT scanner. Arterial phase imaging was performed with fixed delay or bolus triggering, usually between 30 and 35 seconds after injection, and venous phase imaging was performed at 60-70 seconds. All images were reconstructed into thin (0.75-mm slice thickness and 0.5-mm increment) slices. The 3D volume of the pancreas was manually segmented by four trained researchers using commercial segmentation software (Velocity™, Varian Medical Systems Inc.), under the supervision of three abdominal radiologists with 5-35 years of experience. The x, y, and z dimensions of each voxel containing pancreas was determined and summed to calculated the total pancreas volume for each scan. The volume of pancreas was then calculated by counting the number of voxels containing pancreas first and then converting this number into the unit of volume according to the voxel spacing of CT scans.

## RESULTS

### 3D modeling reveals unexpectedly high numbers of spatially separate PanINs in grossly normal human pancreatic tissue

We first sought to use CODA for 3D determination of the size, shape, and number of PanIN lesions in the human pancreas at single-cell resolution (Extended Data 1). From the 38 slabs of grossly normal pancreatic tissue, we examined a mean of 1,288 slides per tissue slab (range 679-1,703) and mean tissue volume of 2.03 cm^3^ (range 0.94-3.62cm^3^) (Supplementary Table 2). Surgical procedures performed included pancreaticoduodenectomy (30/38), distal pancreatectomy (7/38), and total pancreatectomy (1/38). Twelve of these patients underwent surgery to resect a PDAC, and the remaining twenty-six patients underwent resection of other neoplasms not involving the pancreatic ductal system, including pancreatic neuroendocrine tumors, serous cystadenomas, duodenal or ampullary neoplasms, and metastases from other organ sites (Supplementary Table 1). Thirty-six slabs contained exclusively low-grade (LG) PanINs, while two slabs contained PanINs with regions of high-grade (HG) dysplasia.

Across the 38 slabs, 889 spatially separate PanINs were modeled, with a mean number of 23 independent PanINs per slab (range 4-92) (Supplementary Table 2). PanINs contained a mean of 95,021 cells (range 26-7,239,369 cells). Eighty-six percent (764) of PanINs contained less than one hundred thousand cells and 266 (30%) PanINs contained fewer than one thousand cells. (Figures 2E and F). Nine of the 38 tissue slabs analyzed (23%) had at least one large PanIN lesion (greater than one million cells), and the maximum number of such PanINs in a slab was three. The two PanINs containing HG dysplasia were each the largest PanINs by volume in their tissue slabs and contained greater than 200,000 cells each, well above the mean cell count for the all PanINs assessed (Supplementary Table 2). Despite containing thousands of cells on average, the large majority of PanIN lesions (91%) measured <0.5 cm in greatest dimension, consistent with the definitional diameter of PanINs on 2D sections.^5^ We next combined this PanIN cell quantification with the normal ductal cell quantifications in each slab to calculate the percent of epithelial cells in the ductal system that were neoplastic. We found that a mean of 32% of the cells in the ductal system were neoplastic (PanINs) (range 0.2-75%) (Figure 2E). While individual PanINs remain quite small, their cumulative presence is impressive, occupying more than a quarter of the ductal system in most cases.

**Figure 1.**
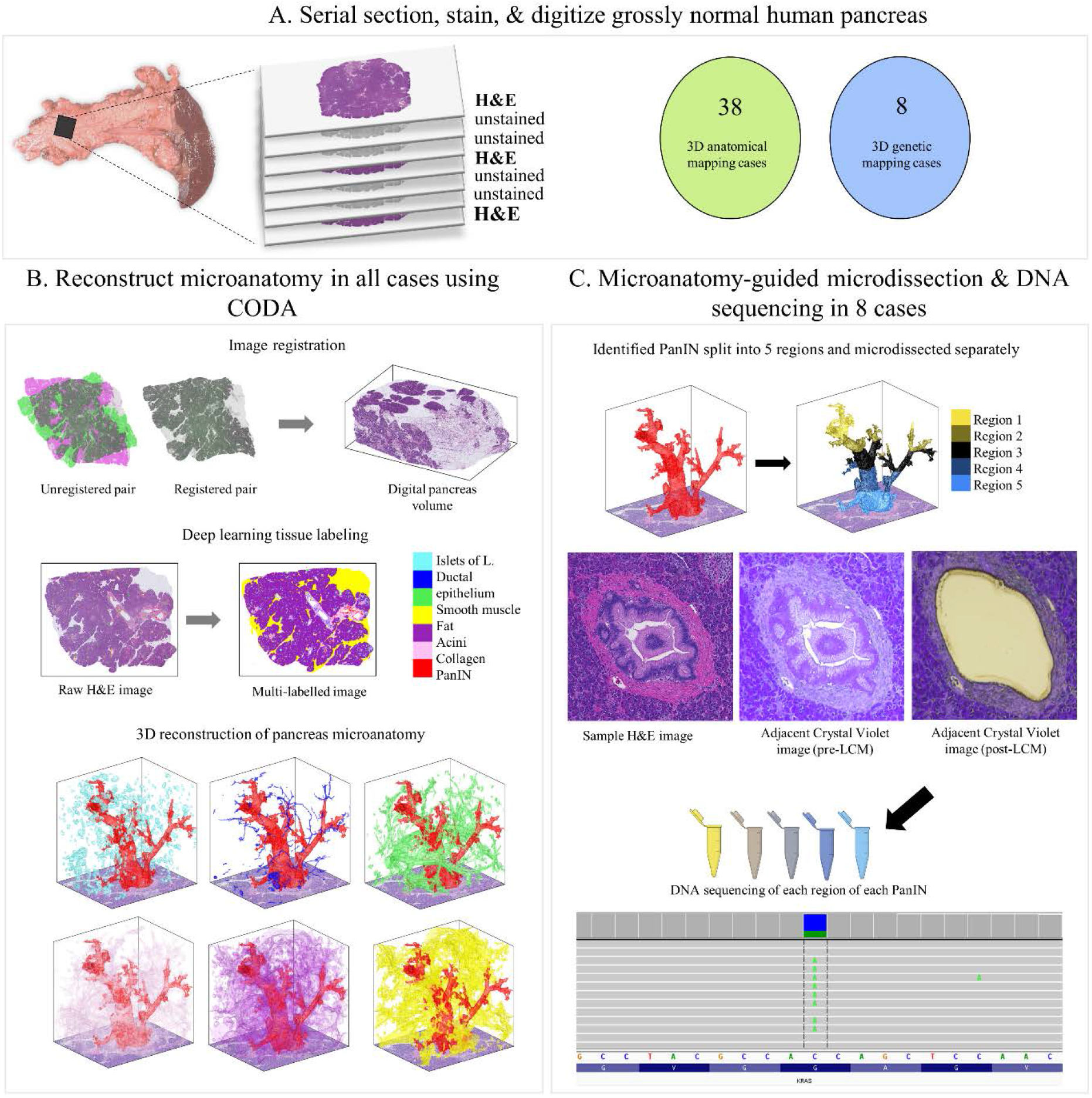
Tissue slab processing for 3D modeling and next generation sequencing. A. Tissue slab processing and sample cohorts. Each grossly normal pancreatic tissue slab was serially sectioned with H&E stains on every third slide for 3D modeling. 38 tissue slabs underwent 3D modeling only. Eight additional slabs underwent 3D modeling and next generation sequencing, with intervening unstained slides used for microdissection to enable 3D-guided genomic analysis. B. 3D reconstruction of tissue slabs utilizing CODA. Registered serial sections allow the creation of a digital pancreas volume. Tissue types are labeled by deep learning, enabling 3D reconstruction of pancreas microanatomy. C. Multi-region microdissection and sequencing of spatially distinct PanINs guided by CODA generated 3D models. Eight tissue slabs underwent multi-region NGS.

**Figure 2.**
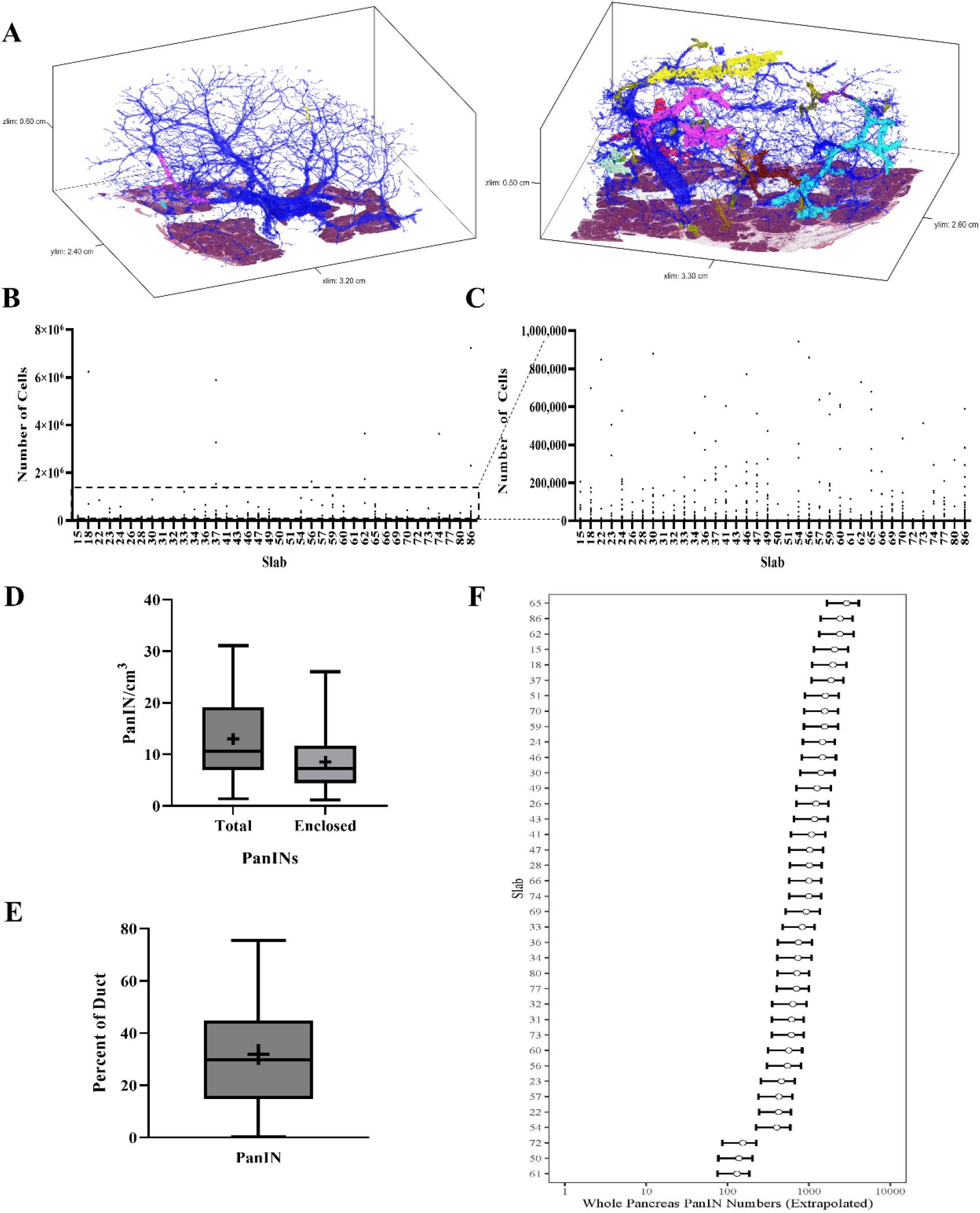
Quantifiable microanatomical features of 3D rendered PanINs. A. Representative 3D models of two human pancreatic tissue slabs. Blue represents normal pancreatic ducts and each spatially distinct PanIN is shown in a unique color. B. PanIN cell counts for all PanINs. The number of cells in each spatially separate PanIN was quantified and shown separately for each tissue slab, as indicated on the x-axis. C. PanIN cell counts for only those PanINs containing less than 1,000,000 cells (subset of data shown in B) D. PanIN burden (PanINs per cm^3^ of analyzed tissue) calculated by total PanINs and those PanINs completely enclosed within the tissue slab. PanIN burden is calculated separately for each tissue slab. + indicates mean. E. Percent of the ductal system affected by PanIN, calculated separately for each tissue slab. + indicates mean. F. Whole pancreas extrapolation of PanIN burden, calculated separately for each analyzed tissue slab.

The overall PanIN burden, calculated as the number of PanINs per slab divided by the tissue volume, was 13 PanINs per cm^3^ (range 4-92 PanINs/cm^3^). Although not statistically significant, this metric was greater in patients with PDAC elsewhere in their pancreas compared to those with non-ductal disease (Extended Data 2A). The trend persisted when comparing the percent of the ductal system affected by PanIN and the number of PanIN cells present in samples from patients with PDAC compared to those with non-ductal disease (Extended Data 2B and 2C). No statistically significant differences in PanIN burden were observed based on patient sex, age, or location of tissue harvest (Extended Data 2D-F). When compared to the proportion of other cell types in the tissue slab, PanIN cell percentage was significantly correlated to percent of normal ductal cells (p=0.0015) and percent of cells in the extracellular matrix (p<0.001) (Extended Data 2 G-H). PanIN cell percentage was significantly inversely correlated to percent of acinar cells (p<0.001), while there was no significant correlation of PanIN cell percentage with percent of fat or islet cells (Extended Data 2 I-L).

The aforementioned calculations likely represent maximum PanIN burdens, as they include all PanINs modeled in the cohort, regardless of their extension beyond the slab analyzed. Of the 889 PanINs modeled, 592 were completely enclosed within the analyzed tissue while 297 were present at a tissue edge (Supplementary Table 2). The 592 completely contained PanINs represent a minimum number of histologically separate PanIN lesions in our cohort. When considering only those 592 completely enclosed PanIN lesions, a mean of 15 independent PanINs per slab were observed (range 4-64) (Figure 2D), leading to a more conservative calculation of mean PanIN burden of 9 PanINs/cm^3^ (range 1-26 PanINs/cm^3^).

To estimate the potential PanIN burden in the entire pancreas, we first determined normal pancreas volumes in 807 kidney donors undergoing CT scans without known pancreatic abnormality (mean and standard deviation of 77.51 cm^3^ and 17.82 cm^3^ in females and 94.18 cm^3^ and 20.74 cm^3^ in males). As the pancreas volumes for the participants in the current 3D tissue modeling study were unknown, we used a sex-specific normal prior distribution for the whole pancreas volume with means and standard deviations indicated above. We obtained a prior predictive distribution for the extrapolated whole pancreas PanIN burden by sampling 10,000 random deviates from the appropriate prior distribution and multiplying by the PanIN burden. Overall, this approach leads to a median extrapolation of 1,021 PanINs in the entire female pancreas and 998 in the entire male pancreas (female range: 139-2,4055, male range: 131-2,902) (Supplementary Table 2) (Figure 2F), in a cohort with a mean age of 65 years old. Even the sample with the lowest PanIN burden led to a whole-pancreas PanIN extrapolation of more than 100 lesions per pancreas, underscoring an unexpectedly large number of PanIN lesions in grossly normal pancreas.

### Combining CODA with next generation sequencing enables 3D genomic analysis of human PanINs

Our 3D reconstruction of grossly normal human pancreas revealed striking multifocality of PanINs within the pancreata of most analyzed patients. However, anatomic analysis alone cannot distinguish whether these spatially unconnected PanIN lesions arose independently or via intraductal spread of a single PanIN. To assess the clonal relationships between and within PanINs, we integrated our 3D anatomic workflow with multi-region DNA sequencing. For this analysis, eight additional large pieces of grossly normal pancreatic tissue were harvested from eight independent surgical pancreatectomy specimens, yielding 37 PanINs for NGS. In this cohort, three patients had a clinical diagnosis of PDAC; the remaining five patients were diagnosed with neoplasms not affecting the ductal system, including neuroendocrine tumors and serous cystadenomas (Supplementary Table 1). To assess both inter-PanIN and intra-PanIN genetic heterogeneity, we micro-dissected each spatially separate PanIN, and neoplastic cells from each PanIN were separately collected across five regions along the z-axis. In addition, when LG and HG dysplasia were present in the same PanIN, these components were isolated separately by microdissection, as was any PDAC identified in the slab. In these eight slabs, we sequenced 99 microdissected samples from 37 PanINs and 5 microdissected samples from PDAC lesions, using a custom targeted panel of 154 well-characterized cancer driver genes, including all major drivers of pancreatic ductal neoplasia (Supplementary Table 3, Supplementary Videos 3-10).

The mean distinct coverage for all targeted sequencing samples was 221X. Of the PanINs in the sequencing cohort, the majority (34/37) contained only LG dysplasia, while three PanINs contained regions of both LG and HG dysplasia. Two slabs in the sequencing cohort contained PDAC within deeper sections of the tissue slab analyzed (Supplementary Table 3). Single nucleotide variants and/or small insertions/deletions were identified in the analyzed PanIN lesions in well-characterized pancreatic driver genes *KRAS* (36/37), *GNAS* (5/37), *RNF43* (2/37), *TP53* (1/37), and *KDM6A* (1/37). Less prevalent somatic mutations were also identified in additional genes on the targeted panel, including *ERBB4* (1/37), *RET* (1/37), *ATRX* (1/37), *STK11* (1/37), *NF1* (2/37), *FLT3* (1/37), and *FGFR3* (2/37). The total number of somatic mutations in the targeted panel identified in each PanIN lesion ranged from 1 to 4, while the two PDACs contained 3 to 5 somatic mutations. *KRAS* was the most commonly mutated gene, with all but 1 PanIN lesion harboring a mutation (36/37 PanINs; 97%), consistent with previous studies^6,13,32^ (Supplementary Table 3). Of these, 6 PanIN lesions harbored *KRAS* p.Q61H mutations (16%), while 30 (81%) were found to have at least one mutation in codon 12 (19 *KRAS* p.G12D; 13 *KRAS* p.G12V; 5 *KRAS* p.G12R; 1 *KRAS* p.G12C) (Extended Data 3). Fifty-four percent of PanIN lesions contained only somatic mutations in *KRAS* in the targeted sequencing analysis, lacking mutations in any other gene in the targeted panel.

To further examine the clonality of single PanIN lesions and the genetic heterogeneity among spatially distinct PanINs in the same slab, we complemented our multi-region targeted sequencing data with whole exome sequencing (WES) and ultra-deep sequencing of *KRAS* hotspots using the Mutation Capsule approach when sufficient DNA remained following targeted sequencing ^25,26^. In total, 57 samples from 24 lesions (20 exclusively LG, 2 with LG and HG dysplasia, 2 PDAC) within the reconstructed tissue blocks were analyzed by WES and deep sequencing of *KRAS* utilizing Mutation Capsule technology (Supplementary Tables 4 and 5). The average distinct WES coverage was 392X (Supplementary table 5). For slabs with a diagnosis of PDAC, we also analyzed a separate PDAC sample, obtained from the archival tissue blocks, to examine the relationship of the analyzed PanIN lesions to the co-occurring PDAC. The number of exomes analyzed per PanIN ranged from 1 to 4 with a mean of 13 somatic mutations per region.

### Histologically distinct PanIN lesions arise from independent genetic events

Because we separately isolated and sequenced DNA from spatially distinct PanIN lesions in each tissue slab, our unique experimental approach allowed for the assessment of inter-lesional genetic differences. All eight pancreatic tissue slabs that underwent 3D genomic analysis contained multiple spatially unconnected PanINs; comparison of somatic mutations between these PanINs can delineate their shared or independent clonal origin. For example, slab 104 had four spatially unconnected PanINs (Figure 3A). Multi-region targeted sequencing analysis revealed that each PanIN in this tissue slab had a distinct *KRAS* hotspot mutation, and no additional mutations in the targeted panel were shared among histologically unconnected PanINs (Figure 3B). Interestingly, this slab included two distinct PanINs containing both LG and HG dysplasia (PanINs C and D). While the LG and HG components within the same PanIN shared the same *KRAS* hotspot mutations, there were discordant *KRAS* hotspot mutations between distinct PanINs.

**Figure 3.**
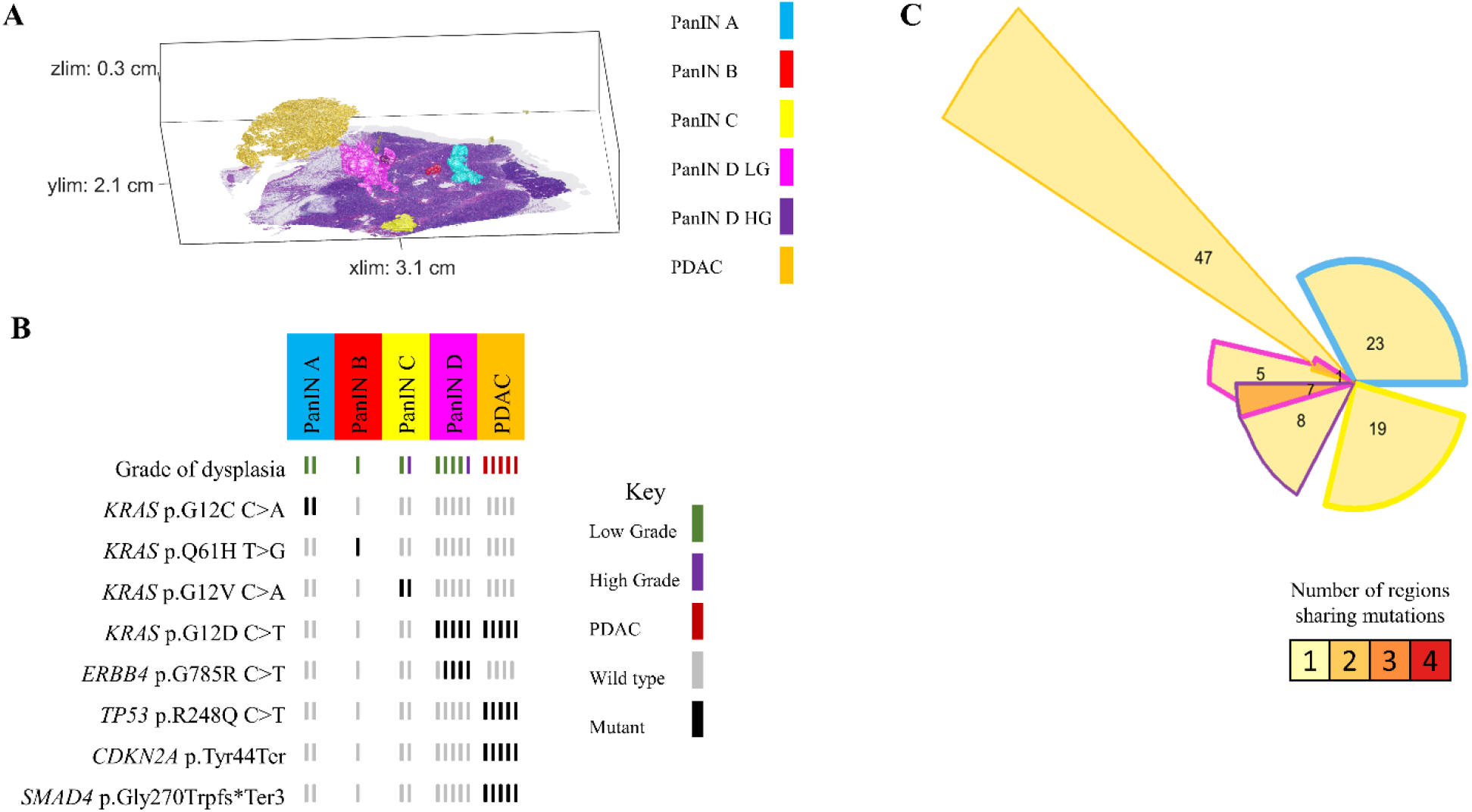
Slab 104 3D model and next generation sequencing results. A. 3D model from grossly normal pancreatic tissue showing multiple spatially distinct PanINs, each represented in unique colors corresponding to key on right. B. Mutation chart with targeted sequencing results. Each column corresponds to a spatially distinct lesion, with lesion colors corresponding to 3D model in A. Each bar represents one region analyzed. Rows display distinct somatic mutations. C. Chow-Ruskey plot summarizing WES results. Each shape represents a group of mutations. Colors outlining each shape correspond to lesion colors in A, with overlapping shapes indicating shared somatic mutations. Numbers within shape indicate the number of mutations present in group.

While these results suggest that the PanINs arose independently, the paucity of driver gene mutations in PanIN lesions make robust conclusions challenging in targeted sequencing experiments. Thus, we compared the somatic mutations identified by WES in seven samples from three spatially unconnected PanINs in slab 104. Although we identified a mean of 19 somatic mutations in each PanIN lesion, no mutations were shared among the spatially unconnected PanINs. In contrast, spatially contiguous regions of the same PanIN lesion shared a mean of 14 somatic mutations, showing the clonal relationship of contiguous regions of the same PanIN lesion. Slab 104 also contained a small focus of PDAC that was analyzed by targeted and exome sequencing. Although this PDAC shared a *KRAS* mutation with LG and HG components of PanIN D in the same tissue block, the PDAC also contained somatic mutations in *CDKN2A*, *TP53*, and *SMAD4* that were absent in the PanIN lesion. Thus, we hypothesized that the PanIN and PDAC in slab 104 were independent neoplasms that shared a common *KRAS* hotspot mutation by chance. This hypothesis was confirmed by the WES data, in which no somatic mutations were shared between the PanIN lesion and the PDAC aside from the *KRAS* hotspot, confirming distinct clonal origins (Figure 3C).

In other analyzed pancreatic tissue slabs, five additional spatially unconnected PanIN lesions shared no genetic alterations with other PanINs in the same slab, indicating an independent clonal origin. Furthermore, six PanIN lesions in this cohort shared only hotspot mutations in *KRAS* with numerous unshared mutations in WES, which we interpret as independent PanINs sharing common hotspot mutations by chance (Figure 5 and Extended Data 8). Despite the presence and proximity of multiple spatially distinct PanINs, they most often arose from independent genetic events. In contrast, three PanINs containing both LG and HG dysplasia shared driver and (when assessed) passenger gene mutations (Figures 3 and 5), indicating the HG PanINs arose from their contiguous LG lesions. Of the three HG PanINs, only one was found to have differences in driver gene mutations with a clonal *TP53* mutation in the HG and subclonal in the LG component. 3D modeling is necessary to delineate spatially distinct lesions and to identify which LG PanIN is contiguous with HG dysplasia.

**Figure 4.**
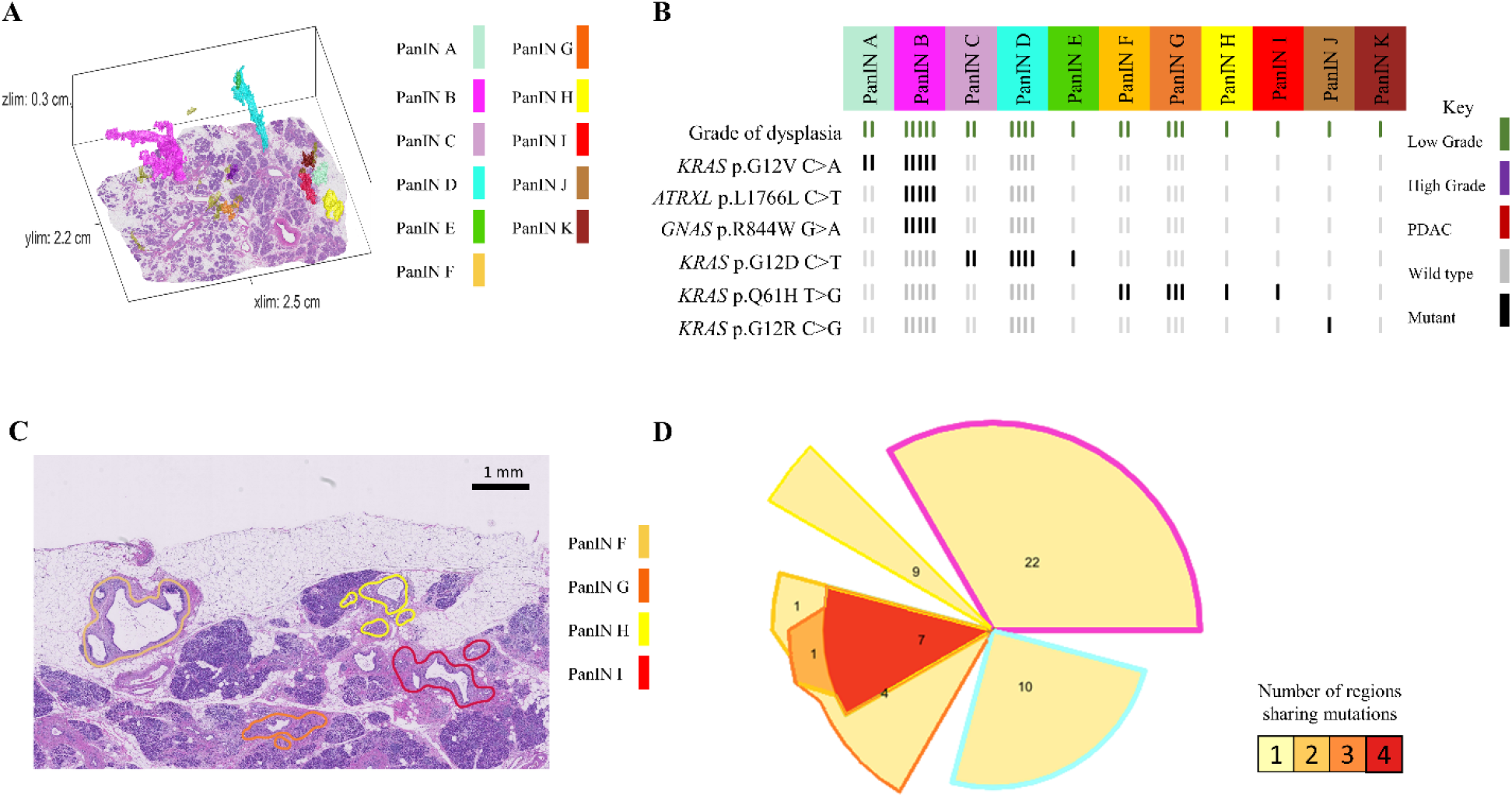
Slab 98 3D model and next generation sequencing results. A. 3D model from grossly normal pancreatic tissue showing multiple spatially distinct PanINs, each represented in unique colors corresponding to key on right. B. Mutation chart with targeted sequencing results. Each column corresponds to a spatially distinct lesion, with lesion colors corresponding to 3D model in A. Each bar represents one region analyzed. Rows display distinct somatic mutations. C. Representative H&E image of PanINs sharing *KRAS* p.Q61H mutations. Each lesion is anatomically distinct from the others, separately by histologically normal pancreatic duct. D. Chow-Ruskey plot summarizing WES results. Each shape represents a group of mutations. Colors outlining each shape correspond to lesion colors in A, with overlapping shapes indicating shared somatic mutations. Numbers within shape indicate the number of mutations present in group.

**Figure 5.**
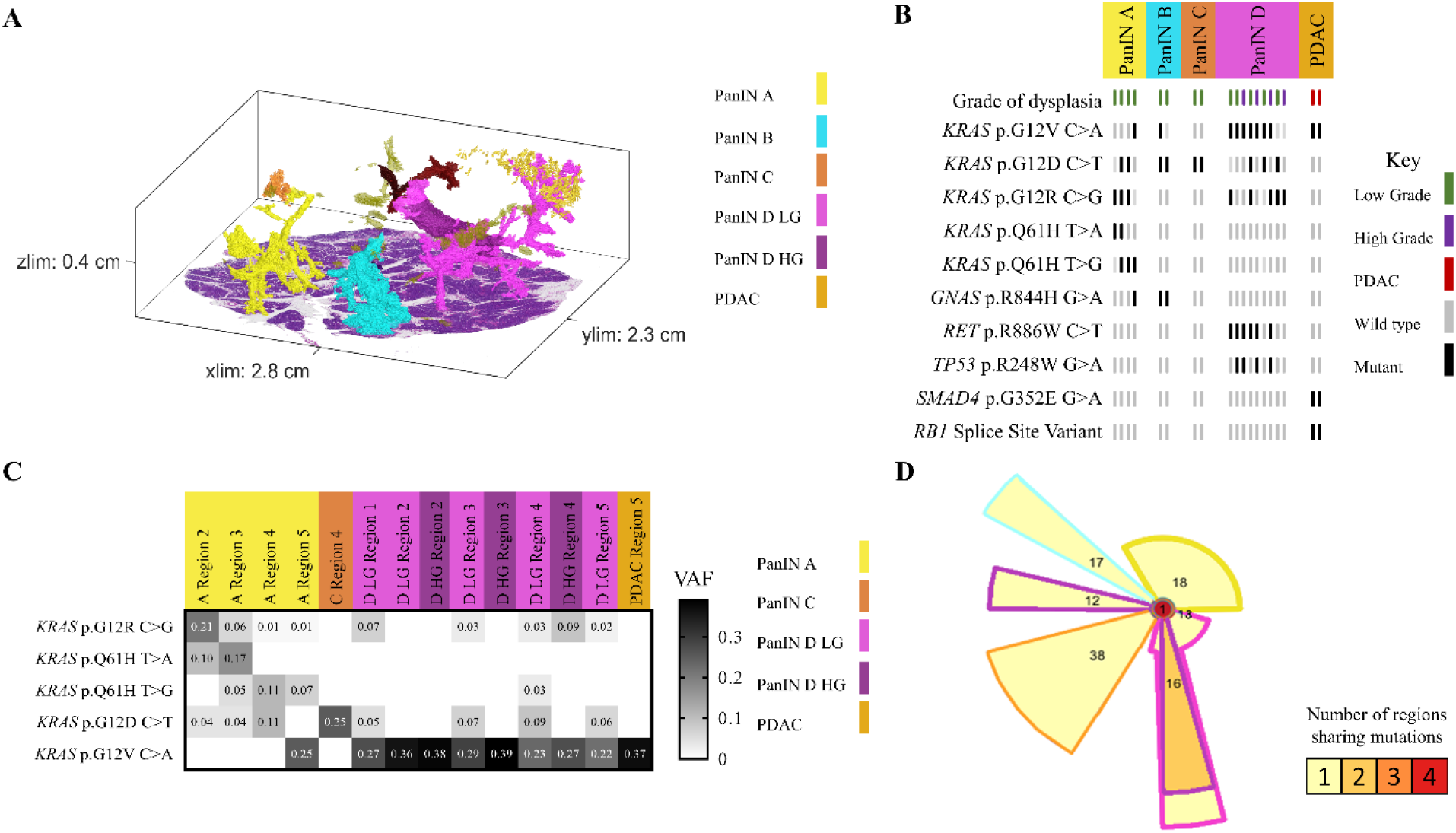
Slab 92 3D model and next generation sequencing results. A. 3D model from grossly normal pancreatic tissue showing multiple spatially distinct PanINs, each represented in unique colors corresponding to key on right. B. Mutation chart with targeted sequencing results. Each column corresponds to a spatially distinct lesion, with lesion colors corresponding to 3D model in A. Each bar represents one region analyzed. Rows display distinct somatic mutations. C. Heatmap of *KRAS* deep sequencing results. Columns represent one lesion with colors corresponding to 3D model in A. Rows display distinct *KRAS* oncogenic hotspot mutations. Variant allele frequency (VAF) indicated within heatmap according to key on right. D. Chow-Ruskey plot summarizing WES results. Each shape represents a group of mutations. Colors outlining each shape correspond to lesion colors in A, with overlapping shapes indicating shared somatic mutations. Numbers within shape indicate the number of mutations present in group.

### Intraductal spread of LG PanIN is rare

In contrast to these common patterns, slab 98 demonstrated the ability of neoplastic cells of LG PanIN lesions to travel short distances via the ductal system, allowing the establishment of spatially unconnected PanIN lesions that share multiple somatic mutations outside of oncogenic hotspots. Of the 11 spatially unconnected PanINs in this slab, four PanINs (PanINs F, G, H, and I) were less than 4mm apart and shared *KRAS* p.Q16H mutations (Figure 4A and 4B). Importantly, PanINs H and I were completely contained within the analyzed tissue block, which excludes connection of the lesions outside of the analyzed tissue. WES was performed on PanINs F, G, and H, while PanIN I lacked sufficient remaining DNA for WES. Interestingly, WES revealed that PanINs F, G, and H shared 7 somatic mutations, though these PanINs were separated by regions of histologically normal non-neoplastic pancreatic duct (Figure 4D). One additional slab, slab 114, contained 2 LG PanINs (PanINs A and B) that shared 7 somatic mutations, including a *KRAS* hotspot mutation. These PanINs were at most 1mm apart but not contiguous, as PanIN B was completely contained within the slab and separated from PanIN A by histologically normal non-neoplastic pancreatic duct (Extended Data 7). Although not common, our 3D genomic analysis confirms the ability of LG PanIN cells to separate from one lesion and establish spatially unconnected but genetically related PanIN lesions nearby.

### Some PanINs may have polyclonal origins

Multiple regions of individual PanIN lesions were separately sequenced, allowing us to assess intra-PanIN genetic heterogeneity. While the majority of PanIN lesions in this cohort harbored one clonal *KRAS* hotspot mutation (81%), 7 PanINs (19%) from 4 slabs (slabs 92, 144, 116, and 117) had multiple *KRAS* hotspot mutations. Slab 92 contained four spatially separated PanIN lesions (Figure 5A). Of these, three PanIN lesions harbored multiple *KRAS* mutations. PanIN A contained 4 distinct *KRAS* hotspot mutations, 2-4 per region, while PanIN B contained 2 unique *KRAS* hotspot mutations. PanIN D contained both LG and HG components. The LG portions had 3 *KRAS* mutations; this heterogeneity was markedly reduced in the HG components. Region 2 of the LG component harbored a *TP53* p.R248W mutation that was clonal in all regions of the HG component (Figure 5B). The intra-PanIN *KRAS* mutational heterogeneity of these PanINs were confirmed by Mutation Capsule analysis, which revealed 5 distinct *KRAS* mutations in PanIN A and 4 distinct *KRAS* mutations in PanIN D (Figure 5C). Because *KRAS* mutations are thought to be very early events in PanIN development, the presence of multiple *KRAS* mutations within a single spatially contiguous PanIN lesion suggests a polyclonal origin for at least a subset of PanIN lesions.

## DISCUSSION

While PanINs are the most common precursor lesions to PDAC, their microscopic size has limited the ability to extensively study their prevalence, spatial landscape, and genetic heterogeneity. Using 3D analyses, we were able to quantify the number of anatomically distinct PanINs in human pancreatic tissues. Remarkably, all tissue slabs assessed contained multiple spatially distinct PanINs, with the number of PanIN lesions identified in each slab ranging from 4 to 92. While the number of anatomically separate PanINs is striking in the tissue we analyzed, our samples account for about 2% of the total pancreas volume; extrapolation of our data to the entire pancreas estimates that a single adult human pancreas can contain hundreds of PanINs. This high PanIN burden is particularly striking considering the relatively low incidence of pancreatic cancer, suggesting that the risk of progression of each individual PanIN is extremely low. These results have important implications for early detection and intervention to prevent pancreatic cancer – future studies should focus on identifying features of PanINs most likely to progress.

The large number of pancreatic precancers in the average pancreas lies in stark contrast to precancers in other organs that are frequently examined, underscoring the need for new paradigms in early detection and intervention in pancreatic neoplasia. For example, colonic adenomas, a well characterized macroscopic precursor to colorectal cancer, are known to be multifocal lesions.^33–36^ However, the number of independent polyps present and prevalence of individuals with multiple lesions are much lower than seen in PanINs in our study. In the absence of inherited cancer predisposition syndromes, patients typically have fewer than five adenomas when the entire colon is examined assessed by colonoscopy or at autopsy.^33,36^ Comprehensive assessment of the burden and multifocality of precancers in other organs will be required to determine whether the high burden of pancreatic precancer is truly unique across organ systems. Our new workflow combining CODA and genetic analysis allows for future extensive 3D analyses of additional microscopic precursor lesions in a variety of organs.

Although previous studies have postulated that multifocal PanIN lesions arise separately, our work definitively demonstrates their independent nature through the development of a technique for integrated 3D modeling and genomic analyses.^6,37–40^ We found that spatially distinct PanINs most often arise from independent genetic events, exemplified by discordant somatic mutations between spatially discrete lesions. Furthermore, we demonstrate that the proximity of LG or HG PanIN to PDAC does not guarantee shared genetic origins – our cohort included 2 examples of PanINs containing HG dysplasia that were genetically distinct from adjacent PDACs, further emphasizing the importance of 3D genomic analyses to ascertain genetic origins. Similar findings have been reported with larger precursor lesions in the pancreas, with a sizable proportion of invasive carcinomas demonstrated to be genetically independent from the co-occurring IPMNs.^17^

An alternative mechanism for developing spatially distinct lesions is the process of intraductal spread. Intraductal spread refers to the ability of neoplastic cells to travel via the pancreatic ductal system to establish spatially distinct but genetically related lesions, and it has been described in pancreatic precursor lesions previously utilizing multi-region NGS. One study described intraductal spread in HG PanINs by demonstrating concordant mutations between lesions appearing to be anatomically distinct on several serial slides.^9^ However, cancerization of the duct by existing PDAC may confound these findings as it mimics HG PanIN histologically,^41^ and connection of the analyzed lesions outside of the examined slides was not definitively excluded. PanINs are complex branching lesions, and, as such, are best studied in 3D. When we performed a rigorous 3D analysis, we found only two unequivocal examples of intraductal spread in our cohort, with adjacent lesions separated by normal ductal epithelium. The intraductal spread of LG PanINs appears to be a rare phenomenon, occurring over short distances.

We also identified significant intra-PanIN genetic heterogeneity, including heterogeneity of driver and passenger gene mutations. This intra-lesional heterogeneity included the presence of multiple *KRAS* mutations in seven PanINs. Because *KRAS* hotspot mutations occur early in the development of PanINs, the presence of multiple *KRAS* mutations in one lesion suggests a polyclonal origin. A polyclonal origin to PanINs is plausible given the previously described polyclonality of IPMNs, with multiple *KRAS* mutations delineating independent clones within the same precancerous cyst.^15^ Furthermore, our results revealed a similar distribution of *KRAS* hotspot mutations in PanIN lesions as have been reported in PDACs (Extended Data 3), suggesting that selection of *KRAS* hotspot mutations occurs at PanIN initiation, not at PanIN progression.^42^

While this study clearly demonstrates the striking multifocality and genetic heterogeneity of PanINs, there are limitations to the cohort assessed. First, grossly normal pancreatic tissue was harvested from patients undergoing resection due to neoplasia, which may not represent truly “normal” pancreas. Still, more than half of the samples were obtained from patients with neoplasms not affecting the ductal system, which may more closely mirror PanIN burden and genetic features in the general population. In addition, due to our sample source of surgically resected pancreata, the age range of the analyzed patients (mean 66, range 45-87) does not reflect the general population. Assessment of PanIN burden in non-diseased pancreata across the age spectrum is an important future direction. While 38 samples were harvested for 3D modeling, combined 3D modeling and genomic analysis was performed on only 8 slabs. Still, in the pancreatic tissue from these 8 patients, multi-region processing yielded one of the most in-depth assessments of somatic mutations in PanINs to date with 99 samples from 37 spatially separate PanINs undergoing NGS. Because LG PanINs are much more common that HG PanINs in randomly sampled pancreata, 34 of these analyzed PanINs contained only LG dysplasia. Although we assessed three HG PanINs and identified clonal expansion of a *TP53* mutation in the HG component of one PanIN, analysis of a dedicated cohort of HG PanINs will be required to make robust conclusions about the molecular alterations driving PanIN progression. Lastly, the genetic origins of seven of the 37 spatially separate PanINs in our sequencing cohort could not be resolved due to a lack of discrete mutations identified by targeted sequencing and insufficient DNA quantities for further WES, underscoring the challenges of genomic analysis of very small precancerous lesions.

As one of the most extensive studies performed with human PanINs to date, we provide new insights into early pancreatic tumorigenesis. 3D modeling alone demonstrated a surprisingly large number of anatomically distinct and morphologically diverse PanINs. Our powerful combination of 3D modeling and genetic analyses revealed that PanINs can be truly multifocal, most often arising from independent clones. While intraductal spread of LG PanINs can occur, it appears to be a rare phenomenon. Rich genetic heterogeneity within individual PanIN lesions was demonstrated by our multi-region sequencing approach, revealing the ability of some PanINs to harbor multiple *KRAS* mutations. The results of this study lay a foundation for future works to determine the underlying mechanisms of PanIN multifocality and for risk stratification of PanIN progression. The current findings describe, for the first time, the spatial and genetic multifocality of human PanINs and underscore the necessity of integrating their 3D microanatomy to accurately resolve genetic origins.

## Acknowledgements

The authors acknowledge the following sources of support: NIH/NCI P50 CA62924; NIH/NCI T32 CA153952; NIH/NCI U54 CA268083; NIH/NCI U54 CA274371; NIH/NCI U54 CA143868; NIH/NCI U54 CA21073; NIH/NIAMS U54 R081774; NIH/NCI U01 CA271273; NIH/NIA U01 AG060903; NIH/NIDDK K08 DK107781; Sol Goldman Pancreatic Cancer Research Center; Lustgarten Foundation; Buffone Family Gastrointestinal Cancer Research Fund; Allegheny Health Network-Johns Hopkins Cancer Research Fund; AACR-Bristol-Myers Squibb Midcareer Female Investigator Grant; Rolfe Pancreatic Cancer Foundation; Joseph C Monastra Foundation; The Gerald O Mann Charitable Foundation (Harriet and Allan Wulfstat, Trustees); Susan Wojcicki and Denis Troper; the Chinese Academy of Medical Sciences (CAMS) Innovation Fund for Medical Sciences (CIFMS) (2021-I2M-1-067 and 2021-1-I2M-018).

## Data Availability Statement

Whole exome sequencing data will be deposited into the European Genome-Phenome Archive (EGA) as allowed by the Institutional Review Board based on patient consent. The 3D rendering software used in this paper is available at the following GitHub page: https://github.com/ashleylk/CODA. Due to their large file size (TB scale per slab), raw tissue data will be available from the corresponding authors upon request.

## Author Contribution Statement

AMB and ALK contributed equally to the work. LDW and DW equally supervised the work. LDW, DW, RHH, and PHW conceived the project. AMB managed the sectioning and scanning of slabs, tissue microdissection, DNA sequencing, and analysis of sequencing results. JMB, LZ, LJ, HC, QS, RR, SG, AID, CGF, SM, CM, JG, XDL, NB, LCC, FL, and NN supported the scanning and sequencing work. ALK managed the computational 3D reconstruction of slabs, anatomical calculations, and visualization of results. MPG, AF, CAP, ACJ, JY, BK, SD, EF, JYH, and PAR supported the 3D reconstruction work. EKF, AY, NJR, EDT, RBS, TCC, YJ, RK, RHH, and PHW oversaw various aspects of the work. AMB, LDW, ALK, and DW created the first draft of the manuscript and figures, which all authors edited and approved.

---

**Extended Data 1.**
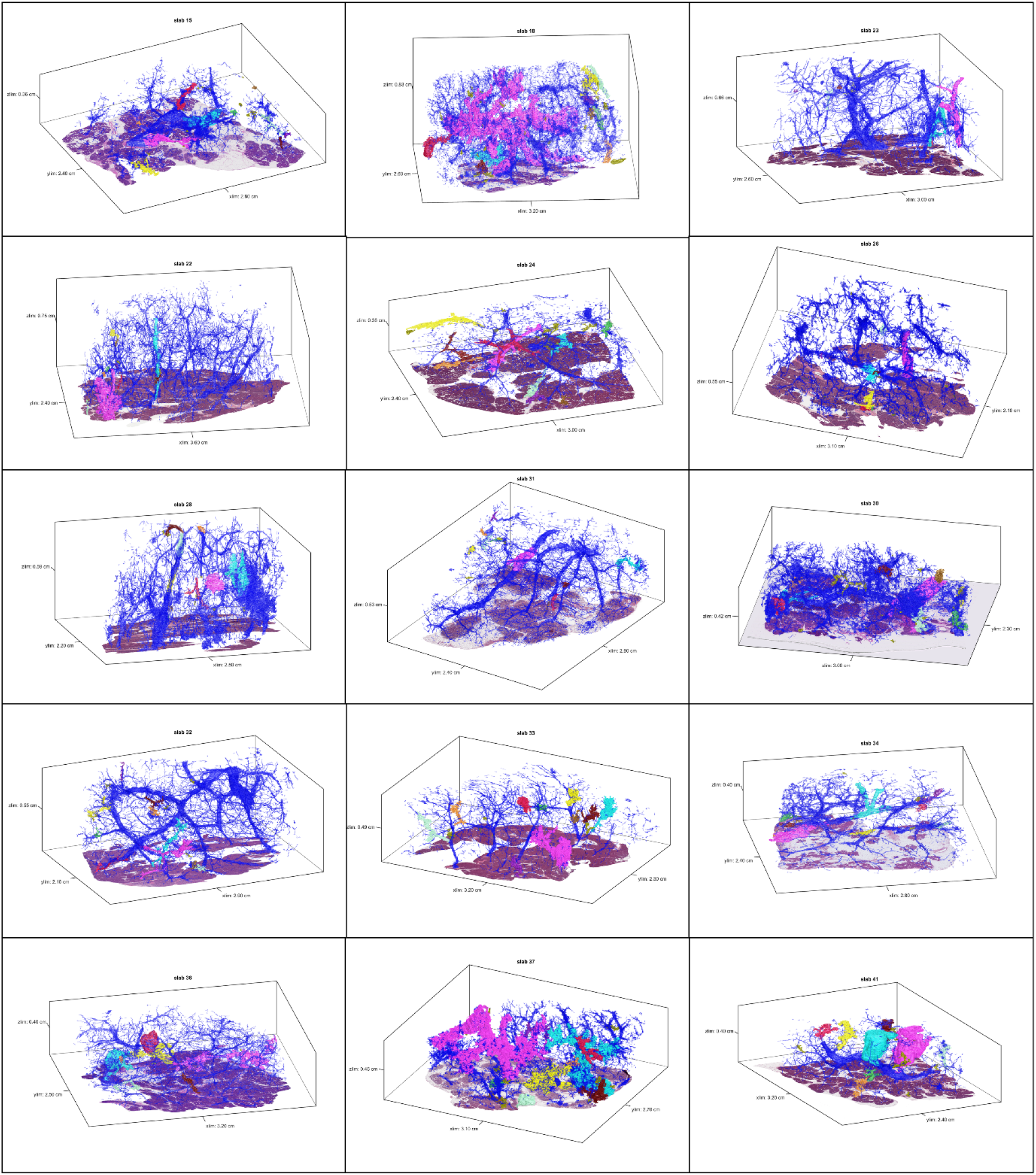
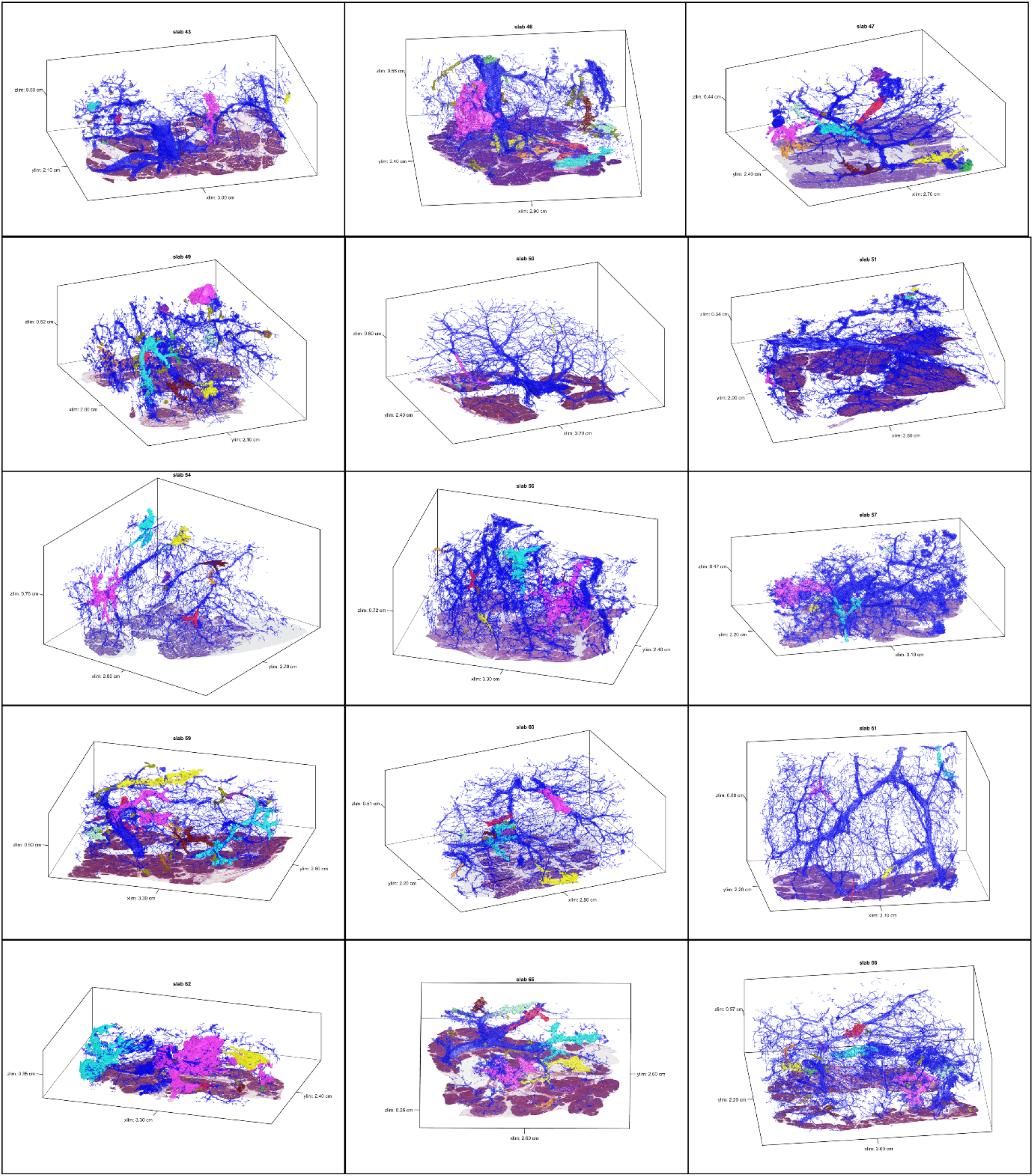
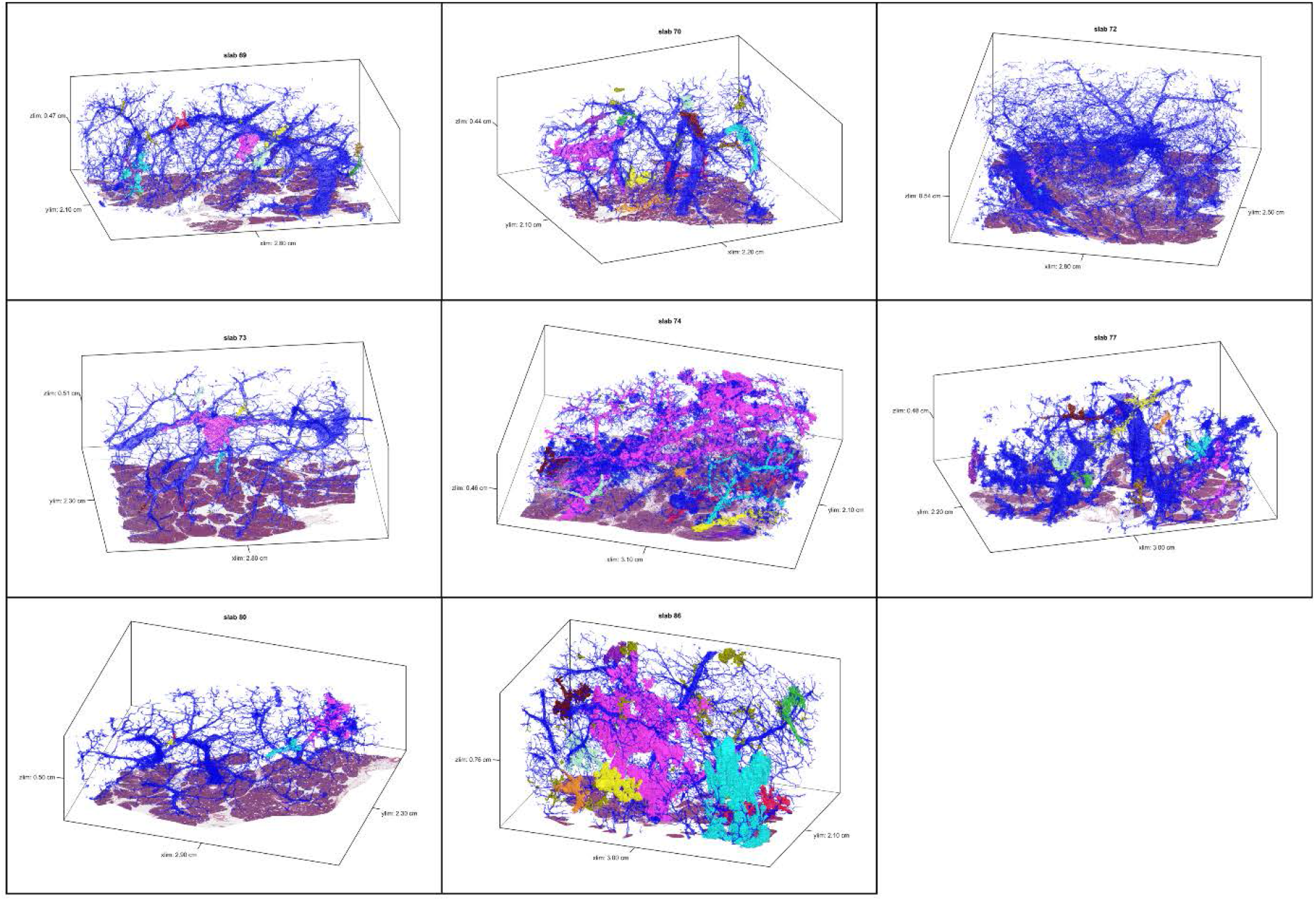
3D models for tissue slabs analyzed. Blue represents normal pancreatic ducts, and spatially separate PanINs are indicated with distinct colors.

**Extended Data 2.**
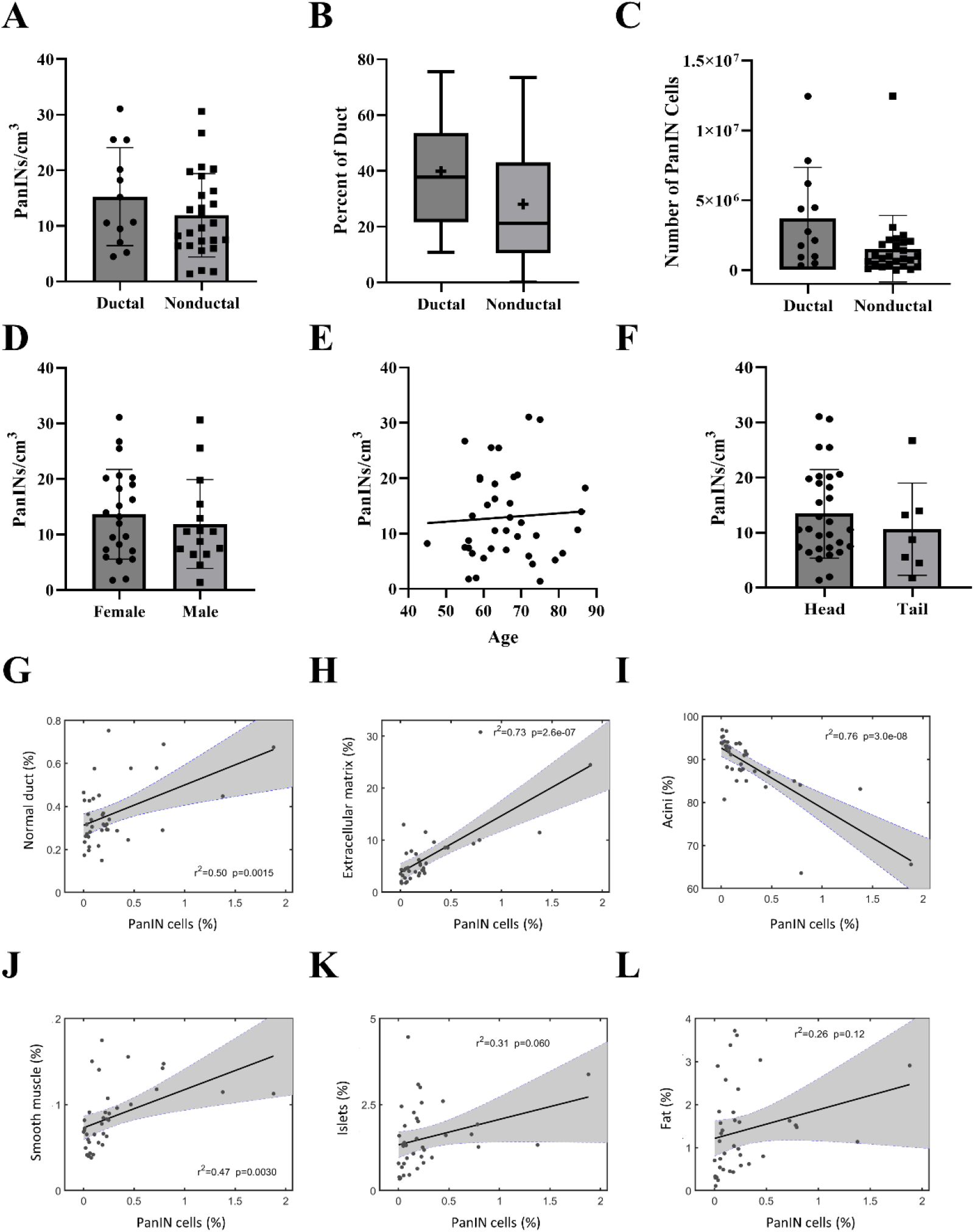
Quantified features from 3D models of human pancreatic tissue slabs. A. PanIN burden stratified by disease type. Spatially separate PanINs per cm^3^ of analyzed tissue was calculated separately for each tissue slab, not statistically significant. B. Percent ductal system affected by PanIN by disease type. Percent of neoplastic ductal cells was calculated separately for each tissue slab, not statistically significant. + indicates mean. C. Number of cells comprising PanINs (calculated separately for each tissue slab), compared by disease type. D. PanIN burden stratified by sex, not statistically significant. E. PanIN burden by age, not statistically significant. F. PanIN burden stratified by location of harvested tissue, not statistically significant. G. Correlation of percent PanIN cells (x-axis) to percent normal ductal cells (y-axis). Each point represents a tissue slab. H. Correlation of percent PanIN cells (x-axis) to percent cells in extracellular matrix (y-axis). p=0.0015. I. Correlation of percent PanIN cells (x-axis) to percent acinar cells (y-axis). p=2.6×10^-07^. J. Correlation of percent PanIN cells (x-axis) to percent smooth muscle cells (y-axis). p=3.0×10^-08^ K. Correlation of percent PanIN cells (x-axis) to percent islet cells (y-axis). Not statistically significant. L. Correlation of percent PanIN cells (x-axis) to percent fat cells (y-axis). Not statistically significant G-L. r^2^ and p values calculated using the correlation coefficient.

**Extended Data 3.**
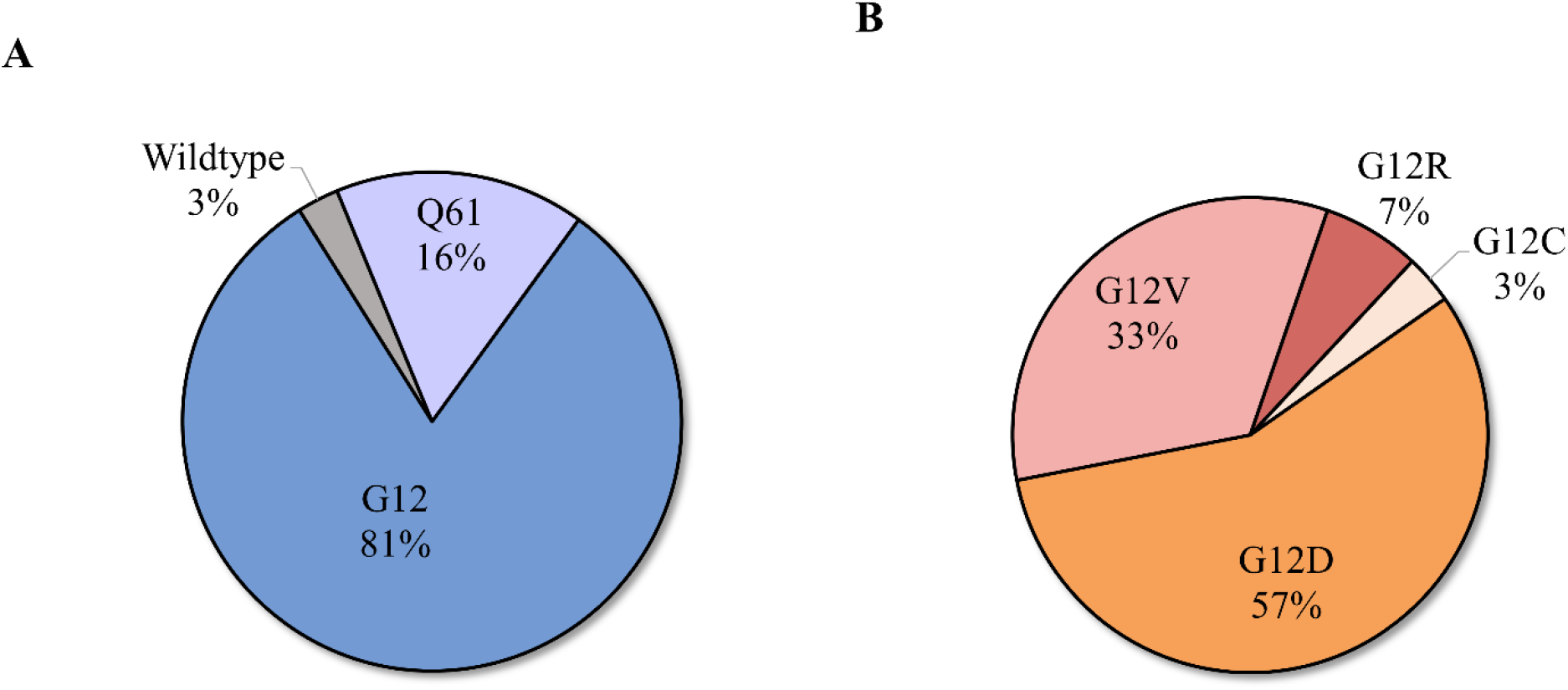
*KRAS* oncogenic hotspot mutations in PanINs assessed in multiregion NGS of eight tissue slabs. A. Missense mutations occur primarily at G12 and at lower frequencies in Q61. B. At the G12 locus, four different amino acid substitutions were found, with p.G12D being the most common.

**Extended Data 4.**
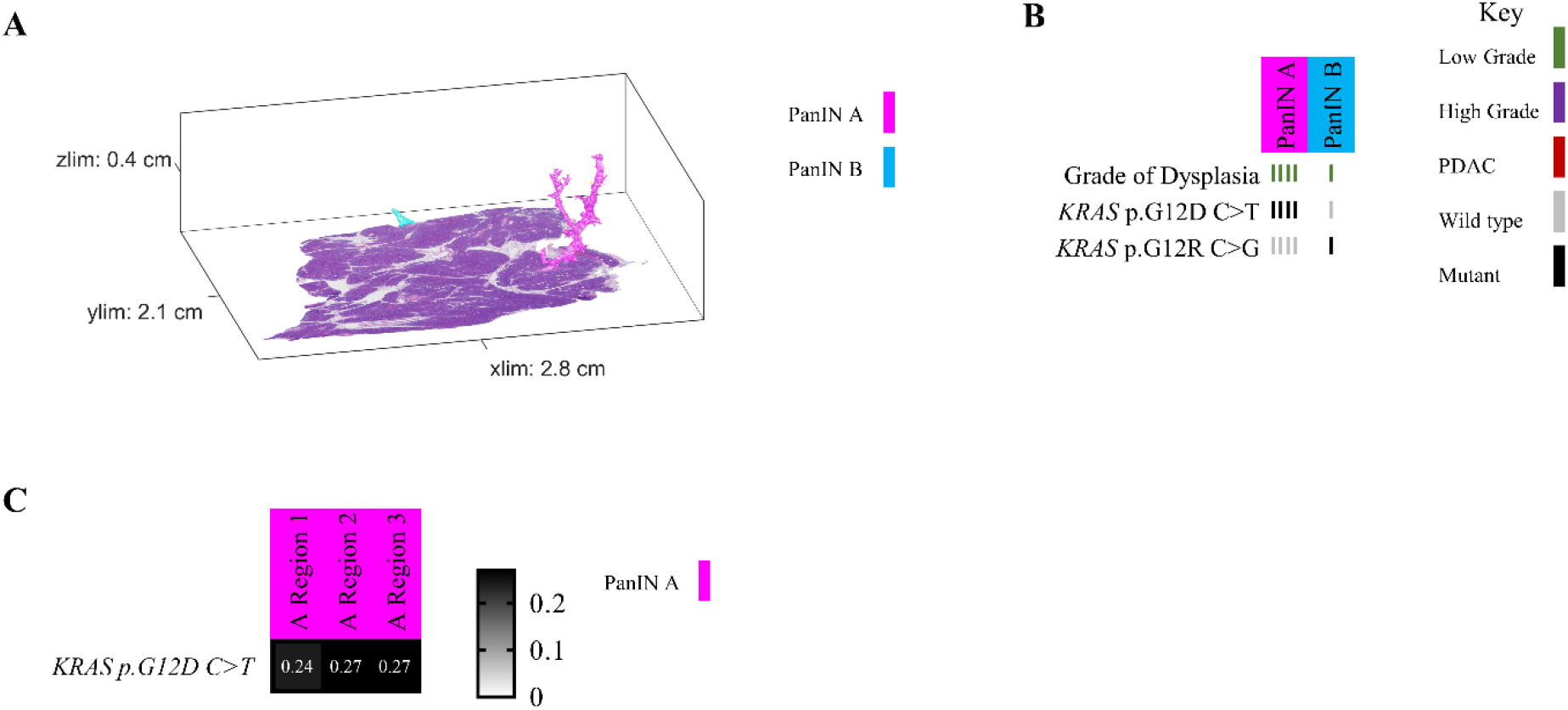
Slab 151 3D model and NGS results. A. 3D model from grossly normal pancreatic tissue showing multiple spatially distinct PanINs, each represented in unique colors corresponding to key on right. B. Mutation chart with targeted sequencing results. Each column corresponds to a spatially distinct lesion. Lesion colors correspond to 3D model in A. Rows display somatic mutations. Each bar represents one region analyzed. Bar color corresponds to key on right. C. Heatmap of *KRAS* deep sequencing results. Columns represent one lesion with colors corresponding to 3D model in A. Rows display *KRAS* oncogenic hotspot mutations. VAF indicated within heatmap and according to key on right.

**Extended Data 5.**
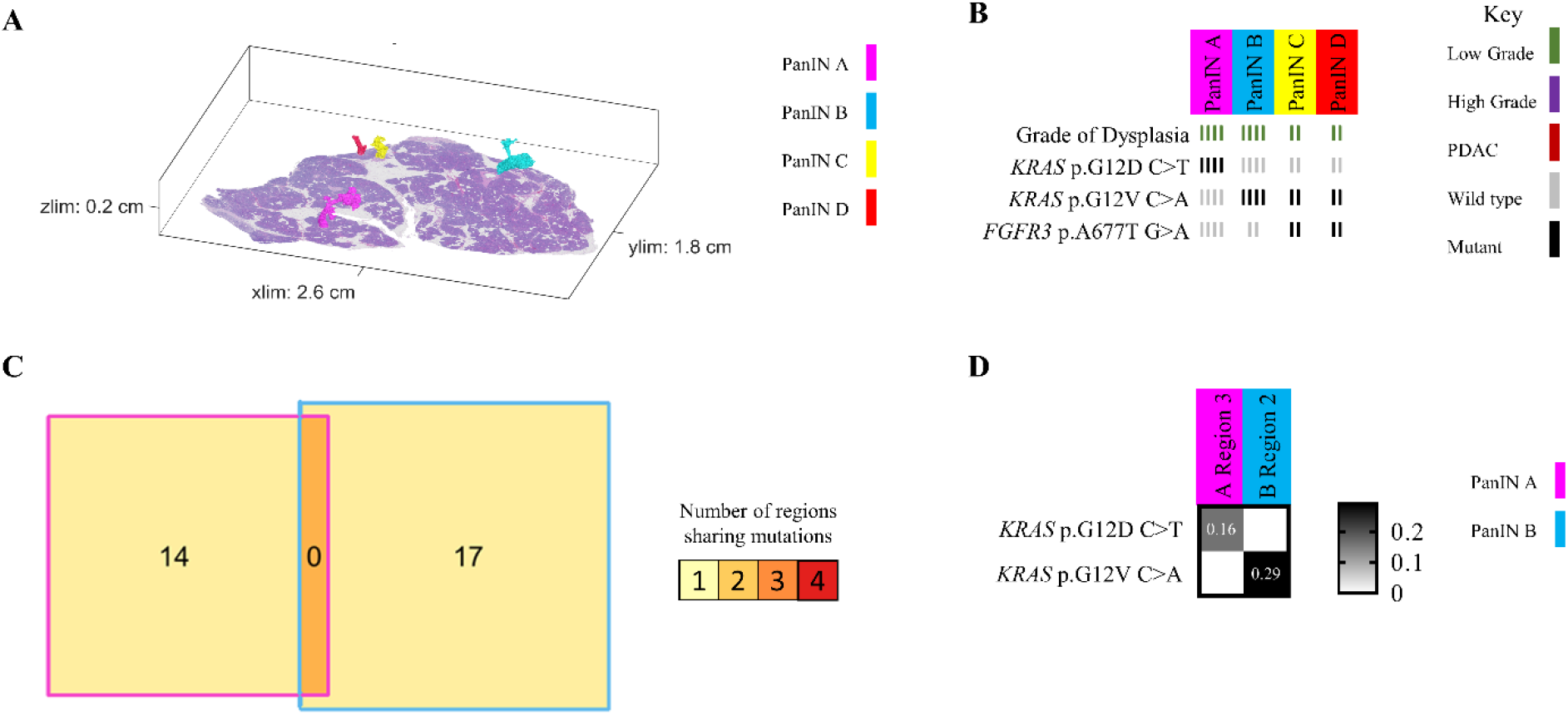
Slab 155 3D model and NGS results. A. 3D model from grossly normal pancreatic tissue showing multiple spatially distinct PanINs, each represented in unique colors corresponding to key on right. B. Mutation chart with targeted sequencing results. Each column corresponds to a spatially distinct lesion. Lesion colors correspond to 3D model in A. Rows display somatic mutations. Each bar represents one region analyzed. Bar color corresponds to key on right. C. Chow-Ruskey plot summarizing WES results. Each shape represents a group of mutations. Colors outlining each shape correspond to lesion colors in A. Overlapping shapes indicate shared somatic. Numbers within shape indicate the number of mutations present in group. D. Heatmap of *KRAS* deep sequencing results. Columns represent one lesion with colors corresponding to 3D model in A. Rows display *KRAS* oncogenic hotspot mutations. VAF indicated within heatmap and according to key on right.

**Extended Data 6.**
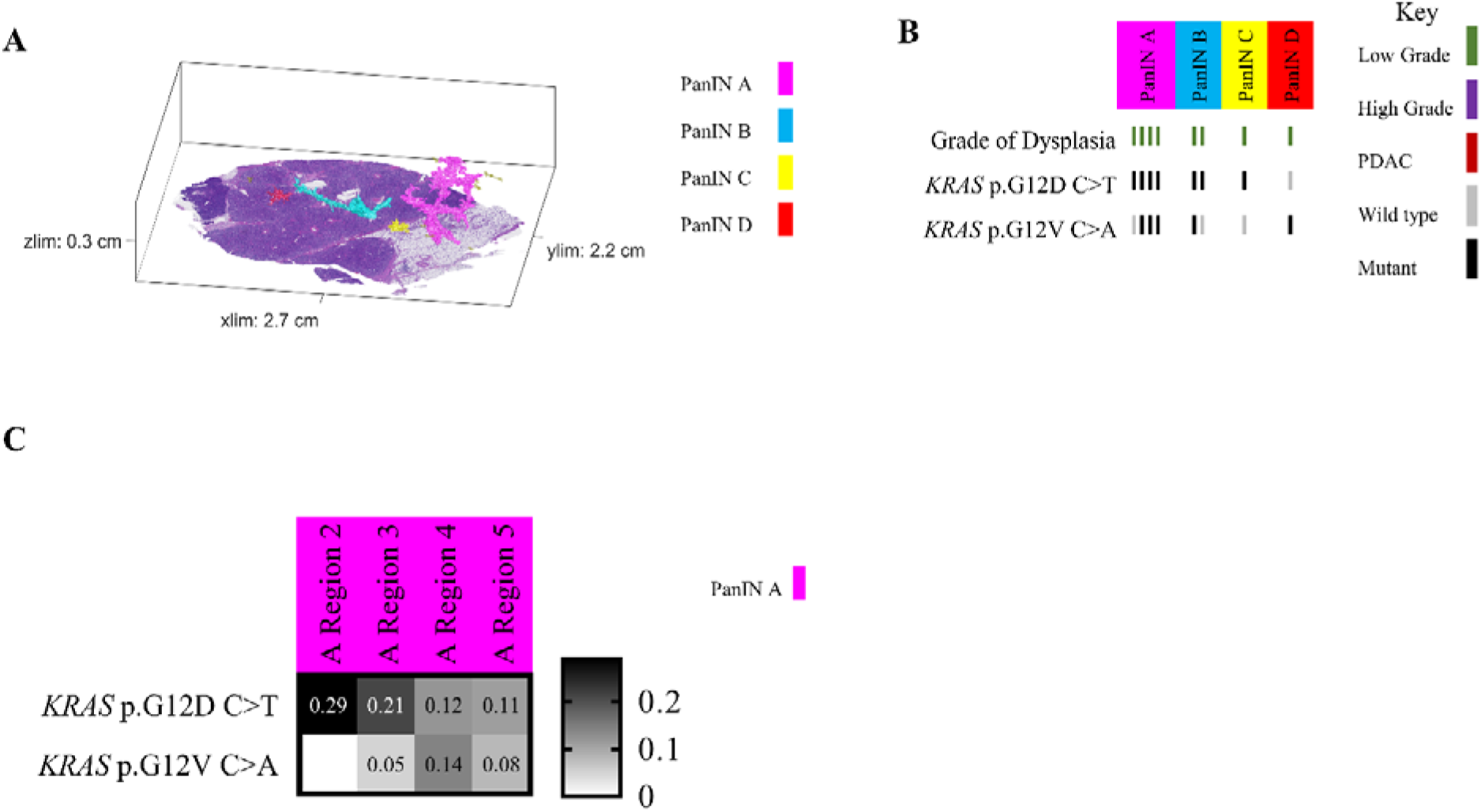
Slab 117 3D model and NGS results. A. 3D model from grossly normal pancreatic tissue showing multiple spatially distinct PanINs, each represented in unique colors corresponding to key on right. B. Mutation chart with targeted sequencing results. Each column corresponds to a spatially distinct lesion. Lesion colors correspond to 3D model in A. Rows display somatic mutations. Each bar represents one region analyzed. Bar color corresponds to key on right. C. Heatmap of *KRAS* deep sequencing results. Columns represent one lesion with colors corresponding to 3D model in A. Rows display *KRAS* oncogenic hotspot mutations. VAF indicated within heatmap and according to key on right.

**Extended Data 7.**
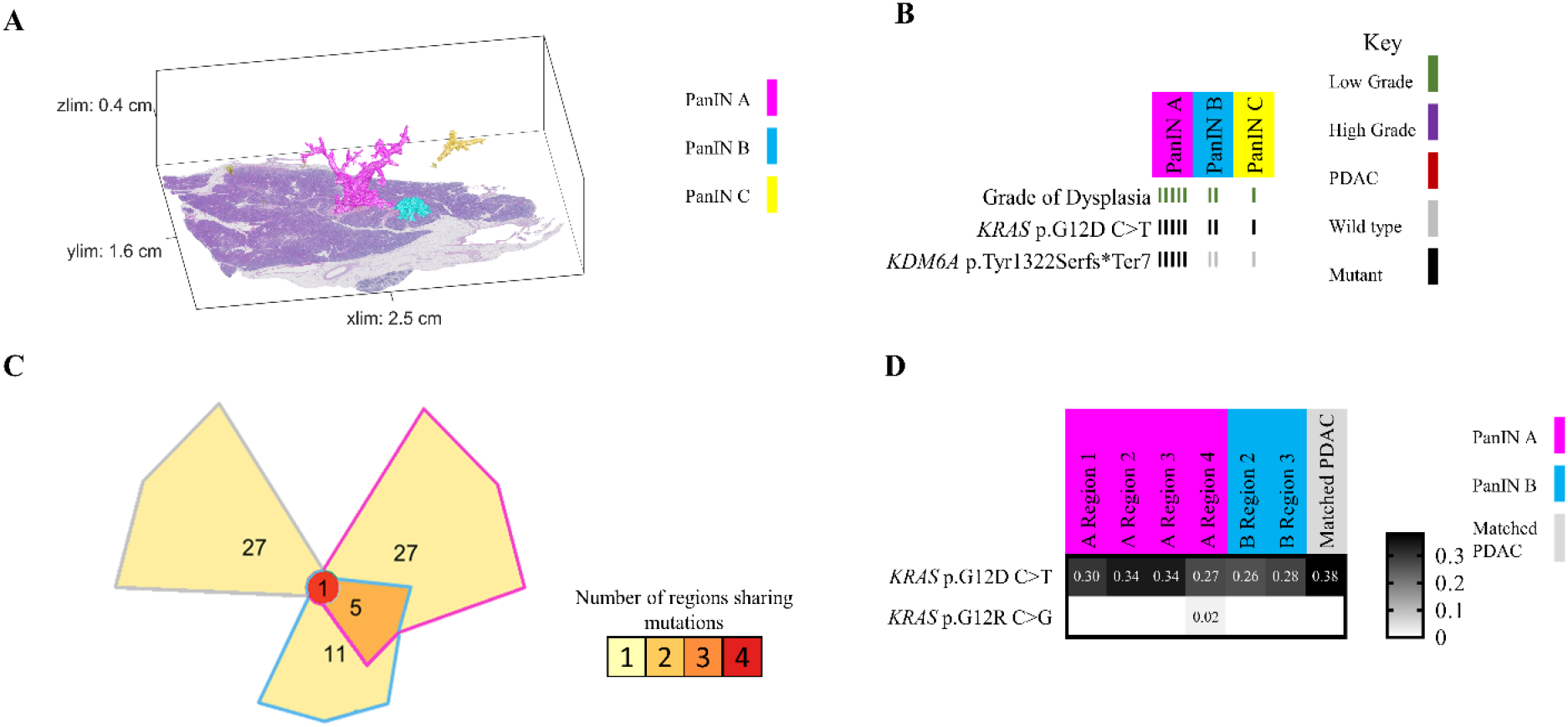
Slab 114 3D model and NGS results. A. 3D model from grossly normal pancreatic tissue showing multiple spatially distinct PanINs, each represented in unique colors corresponding to key on right. B. Mutation chart with targeted sequencing results. Each column corresponds to a spatially distinct lesion. Lesion colors correspond to 3D model in A. Rows display somatic mutations. Each bar represents one region analyzed. Bar color corresponds to key on right. C. Chow-Ruskey plot summarizing WES results. Each shape represents a group of mutations. Colors outlining each shape correspond to lesion colors in A except matched PDAC sample in grey, corresponding to PDAC sequenced from archival clinical block. Overlapping shapes indicate shared somatic. Numbers within shape indicate the number of mutations present in group. D. Heatmap of *KRAS* deep sequencing results. Columns represent one lesion with colors corresponding to 3D model in A except matched PDAC sample in grey, corresponding to PDAC sequenced from archival clinical block. Rows display *KRAS* oncogenic hotspot mutations. VAF indicated within heatmap and according to key on right.

**Extended Data 8.**
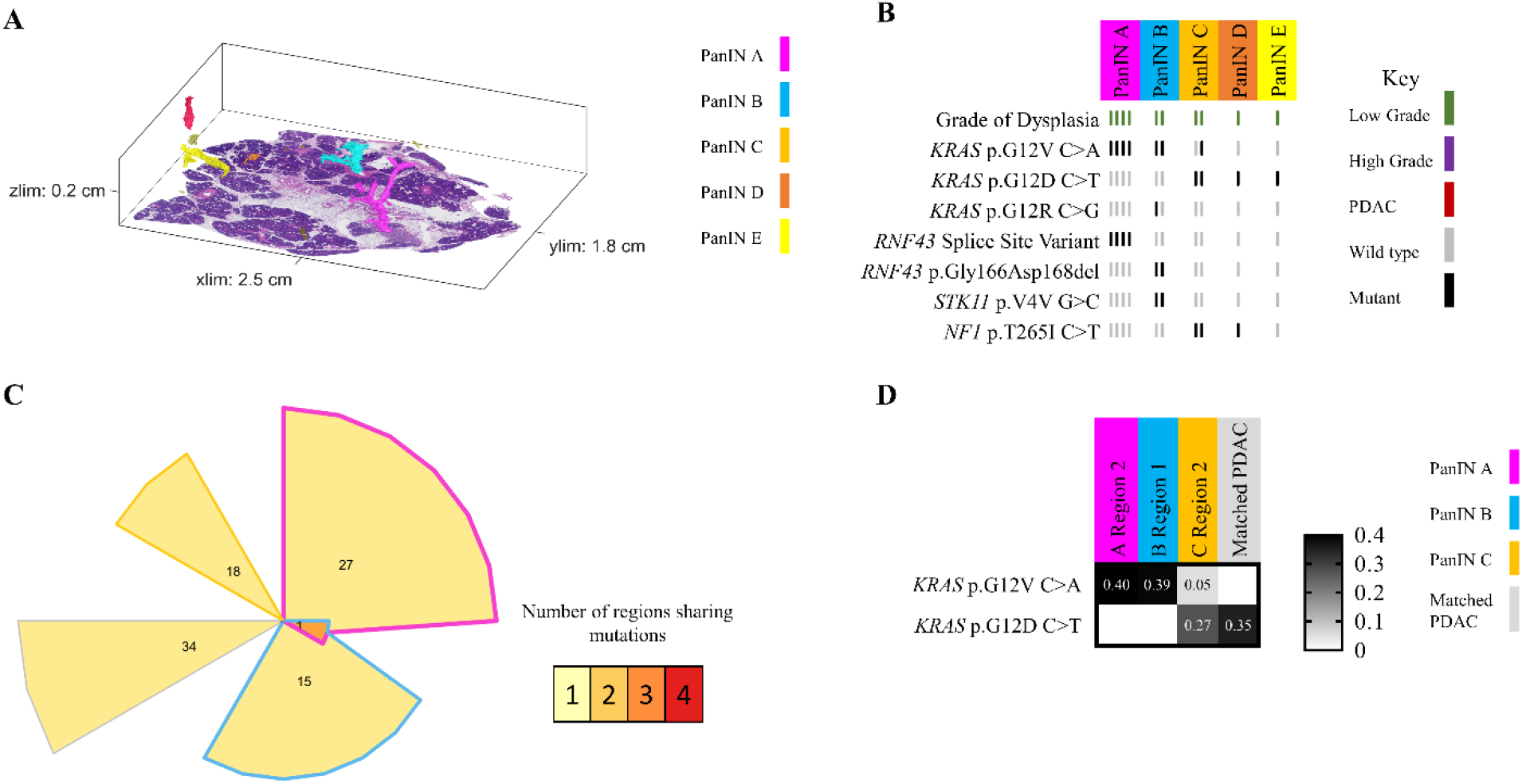
Slab 116 3D model and NGS results. A. 3D model from grossly normal pancreatic tissue showing multiple spatially distinct PanINs, each represented in unique colors corresponding to key on right. B. Mutation chart with targeted sequencing results. Each column corresponds to a spatially distinct lesion. Lesion colors correspond to 3D model in A. Rows display somatic mutations. Each bar represents one region analyzed. Bar color corresponds to key on right. C. Chow-Ruskey plot summarizing WES results. Each shape represents a group of mutations. Colors outlining each shape correspond to lesion colors in A except matched PDAC sample in grey, corresponding to PDAC sequenced from archival clinical block. Overlapping shapes indicate shared somatic. Numbers within shape indicate the number of mutations present in group. D. Heatmap of *KRAS* deep sequencing results. Columns represent one lesion with colors corresponding to 3D model in A except matched PDAC sample in grey, corresponding to PDAC sequenced from archival clinical block. Rows display *KRAS* oncogenic hotspot mutations. VAF indicated within heatmap and according to key on right.

## Supplementary Tables

Supplementary Table 1. Clinical characteristics of samples.

Supplementary Table 2. Quantifications of 3D models.

Supplementary Table 3. Targeted sequencing data and mutation calls.

Supplementary Table 4. *KRAS* deep sequencing data and mutation calls.

Supplementary Table 5. Whole exome sequencing data and mutation calls.

## Supplementary Videos

Supplementary Video 1: Example 3D bounding box for one contiguous PanIN, extracted from the registered 5x magnification image stack. Image stacks were manually viewed using FIJI ImageJ for accuracy of PanIN identification.

Supplementary Video 2: Image sequence of registered, serial H&E sections from slab 92. Spatially separate PanIN were recognized using CODA and annotated on the registered sections for aid in separate microdissection of distinct lesions.

Supplementary Video 3: Three-dimensional rendering of spatially separate PanIN in slab 92. Ten largest PanIN colored distinctly, additional, smaller PanIN colored in olive green.

Supplementary Video 4: Three-dimensional rendering of spatially separate PanIN in slab 98. Ten largest PanIN colored distinctly, additional, smaller PanIN colored in olive green.

Supplementary Video 5: Three-dimensional rendering of spatially separate PanIN in slab 104. Ten largest PanIN colored distinctly, additional, smaller PanIN colored in olive green.

Supplementary Video 6: Three-dimensional rendering of spatially separate PanIN in slab 114. Ten largest PanIN colored distinctly, additional, smaller PanIN colored in olive green.

Supplementary Video 7: Three-dimensional rendering of spatially separate PanIN in slab 116. Ten largest PanIN colored distinctly, additional, smaller PanIN colored in olive green.

Supplementary Video 8: Three-dimensional rendering of spatially separate PanIN in slab 117. Ten largest PanIN colored distinctly, additional, smaller PanIN colored in olive green.

Supplementary Video 9: Three-dimensional rendering of spatially separate PanIN in slab 151. Ten largest PanIN colored distinctly, additional, smaller PanIN colored in olive green.

Supplementary Video 10: Three-dimensional rendering of spatially separate PanIN in slab 155. Ten largest PanIN colored distinctly, additional, smaller PanIN colored in olive green.

